# Lossless end-to-end transport of small molecules through micron-length DNA nanochannels

**DOI:** 10.1101/2022.04.13.488239

**Authors:** Yi Li, Christopher Maffeo, Himanshu Joshi, Aleksei Aksimentiev, Brice Ménard, Rebecca Schulman

**Affiliations:** Chemical and Biomolecular Engineering, Johns Hopkins University, Baltimore, Maryland 21218, United States; Physics and Astronomy, Johns Hopkins University, Baltimore, Maryland 21218, United States; Computer Science, Johns Hopkins University, Baltimore, Maryland 21218, United States; Department of Physics, University of Illinois Urbana-Champaign, Urbana, IL 61801, United States

## Abstract

Designed and engineered protein and DNA nanopores can sense and characterize single molecules and control transmembrane transport of molecular species. However, designed biomolecular pores are less than 100 nm in length and are used primarily for transport across lipid membranes. Nanochannels that span longer distances could be used as conduits for molecules between non-adjacent compartments or cells. Here, we design microns-long, 7 nm diameter DNA nanochannels that small molecules can traverse according to the laws of continuum diffusion. Binding DNA origami caps to channel ends eliminates transport and demonstrates that molecules diffuse from one channel end to the other rather than permeating through channel walls. These micron-length nanochannels can also grow, form interconnects, and interface with living cells. This work thus shows how to construct multifunctional, dynamic agents that control molecular transport, opening new ways of studying intercellular signaling and modulating molecular transport between synthetic and living cells.

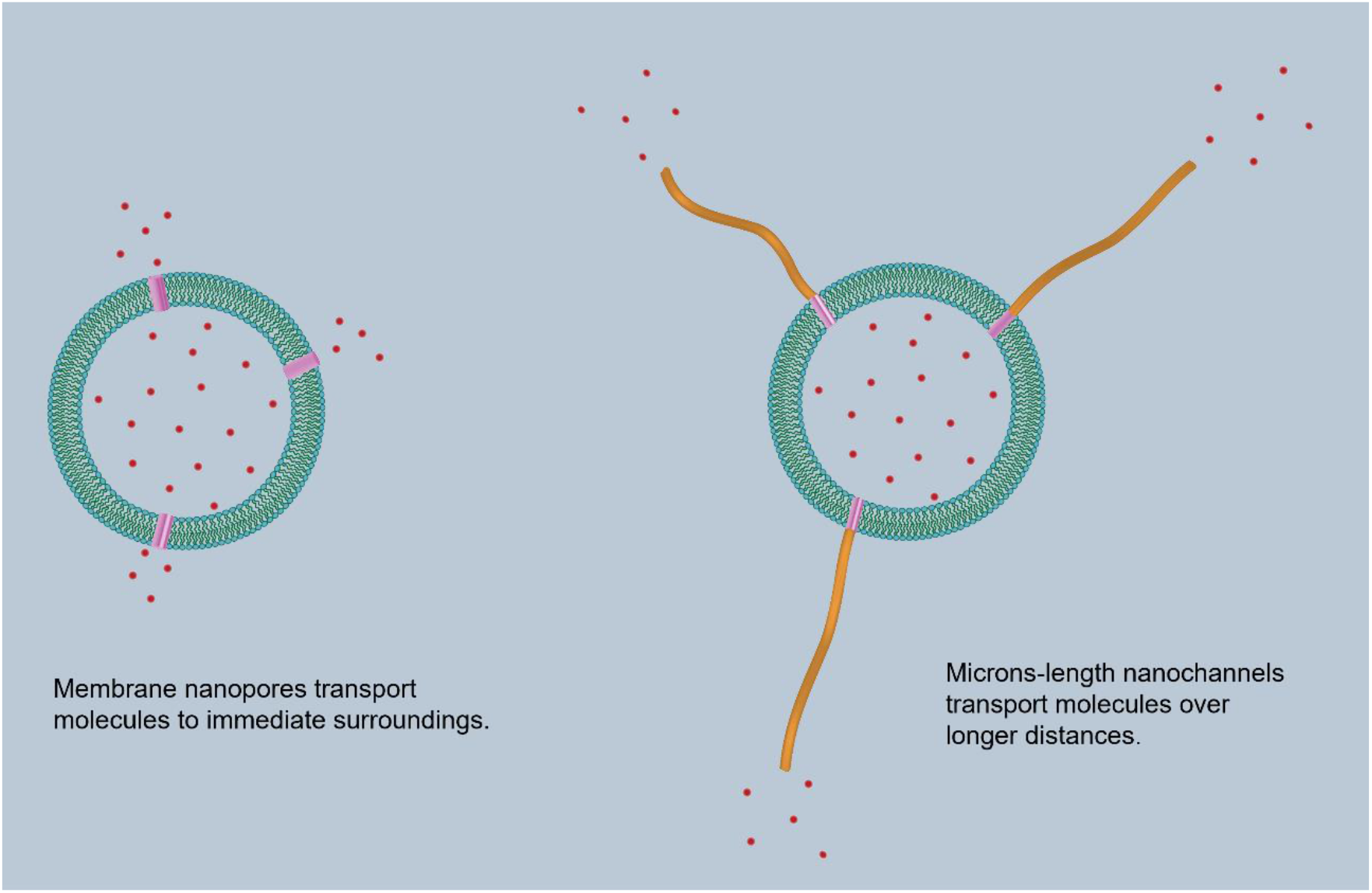

## Introduction

Nanoscale channels are fundamental mechanisms for directing transport across membranes in living systems. Synthetic nanopores and nanochannels that mimic the selectivity^1–3^ and gating^4,5^ functions of biological membrane channels have become powerful tools in biosensing^6–8^ and drug delivery^9,10^. Biomimetic channels assembled from different materials, such as peptides^11,12^, polymers^13^, DNA^14^, metal-organic complexes^15,16^, and carbon nanotubes^17^, can mediate cross-membrane transport of a range of ions and molecules. Most of these biomimetic channels have diameters below 2 nm and focus on studying the transport of ion species.

Self-assembled DNA-based synthetic nanopores, particularly those built from scaffolded DNA origami, benefit from the extensive design space of DNA as building materials. DNA nanopores with inner diameters over 3 nm have been shown to mediate the passage of large biomolecules, such as double-stranded DNA and proteins, across lipid bilayer membranes.^18–22^ Moreover, the highly predictable DNA interactions are the basis for the creation of programmable DNA nanopores that initiate transport only in the presence of specific chemical or spatial cues.^20,23^ Advancements in DNA modifications with various chemical groups and aptamers and control over pore geometries have further enabled specificities in the selective transport of solute species across the DNA nanopores.^21,24^ In addition to the potential applications in biosensing and drug delivery,^25^ DNA nanopores can serve essential functions as transporters in systems of synthetic cells^26^: The lack of specificity of many types of membrane pores and the lack of reliable transport mechanisms between synthetic cells are central challenges in reconstituting complex signaling systems in synthetic cells.^27^ The programmability and specificity of DNA nanopores would allow delivery of target genetic and signaling materials to make efficient communications possible.

However, existing large-diameter DNA nanopores have lengths below 100 nm and act as cross-membrane transporters.^28^ Short nanopores only allow transport within immediate environments. Longer DNA-based channel structures, once built, can be used to establish connections between separated cells or compartments for communications and molecular exchange. Furthermore, such channels can enable the development of complex transport networks that connect multiple components in specific ways. Nevertheless, such longer DNA channels have not been reported before, mainly due to the size limitation^29^ of the 6-helix bundle stem and DNA origami-based designs that are typically used for building DNA nanopores. It also remains unknown whether the end-to-end passage of molecules can occur after such longer DNA channels are built.

Here, we demonstrate bulk transport of molecules over distances through microns-long DNA-based nanochannels and investigate the transport phenomena within them (Fig. 1a). DNA nanochannels that span up to several microns in length are self-assembled from DAE-E double-crossover DNA tiles, on top of DNA origami pores of an approximately 7-nm inner diameter. Observations that fluorescent dyes move across lipid membranes confirm that molecular transport can occur through both DNA pores and nanochannels. Further analysis of the kinetics of these processes suggests a continuum diffusion description of the transport within nanopores. By showing a DNA origami cap that closes the channel opening prevents channel-mediated transport, we demonstrate specific gating of the nanochannel and that the transport is predominantly end-to-end rather than across channel walls.

**Figure 1.**
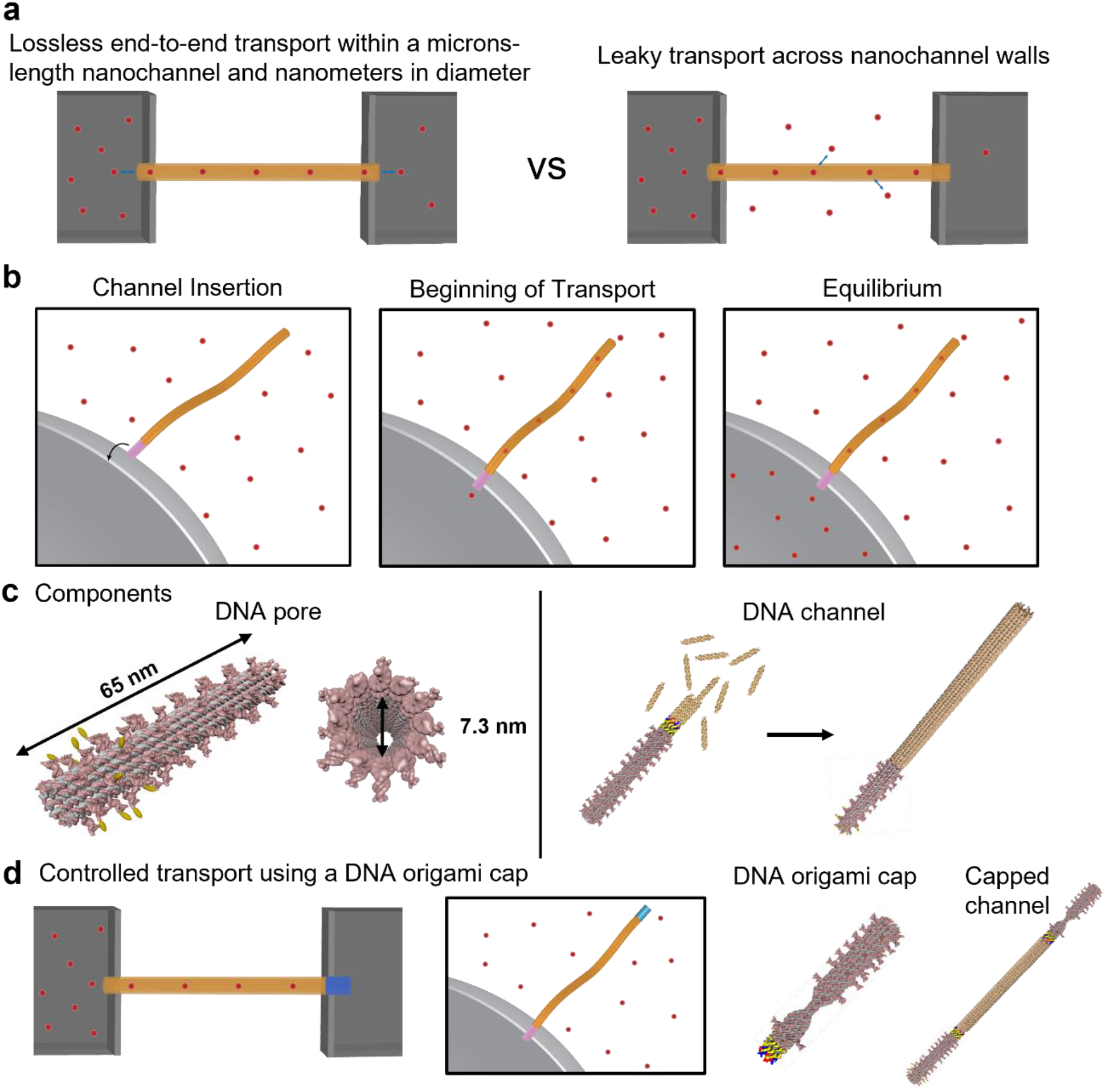
Scheme for studying the transport of small molecules through micron-length self-assembled DNA nanochannels. (**a**) End-to-end transport of small molecules through micron-length channels could allow for controlled transport between channel endpoints. Diffusion across the channel walls leads to transport loss. (**b**) The assay used to characterize the rate of transport through a micron-long channel. When the channel attached to a pore enters a lipid membrane of a giant unilamellar vesicle (GUV), fluorescent dyes diffuse into the vesicle because of the concentration gradient across the membrane until equilibration of the concentration of fluorescent dyes inside and outside the vesicle. (**c**) A DNA origami nanostructure functionalized on one end with hydrophobic moieties serves as a transmembrane pore. DNA tiles hybridize to sticky ends on the DNA pore’s opposite end to self-assemble a DNA nanotube hundreds of nanometers to microns in length that serves as a nanofluidic channel. (**d**) A cap that binds to and plugs the opening of the nanotube channel would prevent transport through the channel end into the vesicle but not transport across channel walls. A DNA channel cap with a designed constriction binds to the nanotube channel end via complementary DNA sticky ends.

The designed DNA nanochannels thus mediate lossless transport from one end of a barrel to the other over distances ranging from tens of nanometers to microns, and this transport can be completely halted by the programmable binding of a DNA cap. The rates of transport in these channels are consistent with those in bulk solutions, allowing for precise design of flow rates. These advances suggest a toolkit of self-assembling elements for building nanoscale fluid transport networks to connect compartmentalized structures separated in space. Such transport networks have great potential for studying intercellular communications and building artificial multicellular structures.

## Results

### Nanochannel Design and Structural Characterization

To measure the rate of small-molecule transport through self-assembled DNA nanochannels, we first developed a DNA pore that could be inserted into lipid bilayer membranes. Molecules diffuse through the pore into a lipid vesicle, whose concentration is monitored over time (Fig. 1b). The DNA pore serves as both a submicron-length cross-membrane channel and as a template for the growth of micron-length nanochannels.

We constructed such a DNA pore by modifying a 12-helix scaffolded DNA origami cylinder of length 63.1 ± 1.8 nm, a nanotube seed^30^, that can act as a template for DNA nanotube growth (Fig. 1c left, Supplementary Note 1). We added 15-nt extensions to 12 of the staple strands near one end of the structure to which 12 cholesterol-conjugated ssDNA strands could hybridize. The diameter of the pore was measured to be 7.3 ± 0.4 nm at the end of equilibrium coarse-grained MD simulations performed using mrDNA^31^ (Supplementary Fig. 4). Electron micrographs of >100 pores (Fig. 2a, Supplementary Fig. 6) showed a mean length of 61.1 ± 0.5 nm, consistent with prior measurements via atomic force microscopy^30^. Cholesterol-modified pores could spontaneously insert into the lipid bilayer membranes of small unilamellar vesicles (SUVs) at orientations generally perpendicular to the membranes (Fig. 2c).

**Figure 2.**
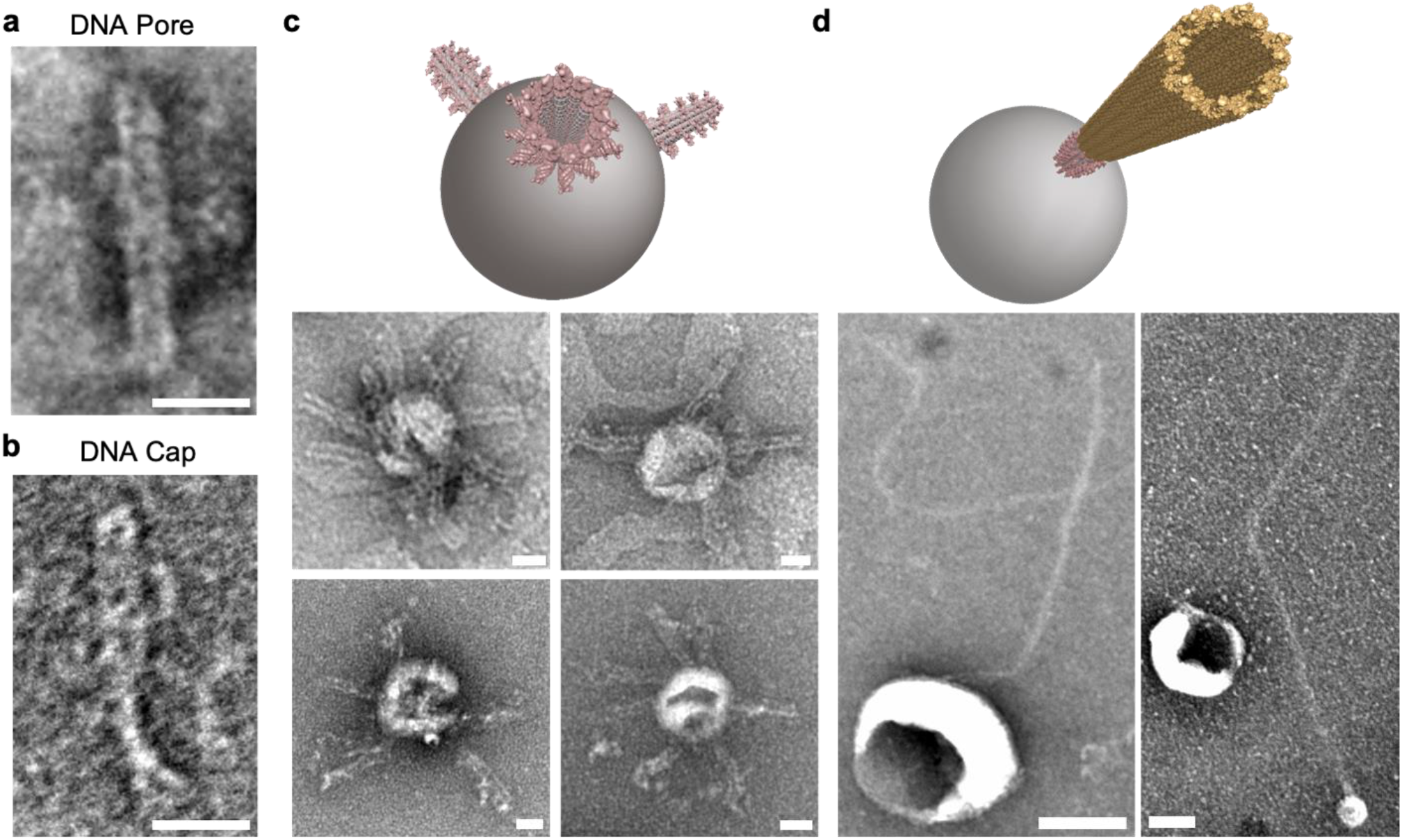
TEM characterization of DNA pores and DNA nanotubes with small unilamellar vesicles (SUVs). (**a**) Transmission electron micrograph of a DNA origami pore without DNA-cholesterol conjugates (Supplementary Note 4). (**b**) Transmission electron micrograph of a DNA channel cap. (**c**) Schematic and images of cholesterol-modified DNA origami pores interacting with SUVs. (**d**) Schematic and TEM images of cholesterol-modified DNA interacting with SUVs. Scale bars 20 nm (**a**-**c**), 100 nm (**d**).

We assembled longer nanochannels by growing nanotubes from the DNA pores hundreds of nanometers to several microns in length (Supplementary Note 2, 3) during an isothermal incubation at 37 °C (Fig. 1c right). We then attached cholesterol-DNA conjugates to the pores. These channels could also interact with lipid bilayer membranes after being mixed with SUVs (Fig. 2d).

We designed a scaffolded DNA origami cap that binds to the open end of a DNA pore or a nanotube via sticky-end hybridization by modifying a rigid nanotube cap^32^ (Figure 1d, Supplementary Fig. 3). Pore-assisted small molecule transport through capped channels could occur only if the small molecule could readily diffuse transport across channel walls, *i.e*., through interhelix gaps or structural defects. We thus hypothesized that by comparing rates of pore-mediated molecular transport across capped and uncapped channels, we could determine whether nanochannels could direct the end-to-end transport of small molecules across their full lengths without loss through channel walls.

### Transport rates through DNA pores suggest a continuum diffusion model

Controlling small molecule transport through microns-long channels would be difficult if transport was mediated by specific molecule-surface interactions; molecule aggregation might also form clumps obstructing the channel. We therefore first sought to understand whether molecular transport occurs within the DNA channels by measuring rates of small molecule transport through the pores^33,34^ and comparing these rates to the predictions of macroscopic diffusion models.

We tracked the flux of a fluorescent dye, carboxytetramethylrhodamine (TAMRA), through DNA pores into giant unilamellar vesicles (GUVs). TAMRA was selected for its small size, neutral charge, and low membrane permeability.^35^ GUVs were immersed in a TAMRA solution containing 1 nM DNA pores and immobilized to a glass coverslip via biotin-streptavidin linkages. The rates of TAMRA influx into GUVs over a 3.2-hour duration were observed using time-lapse confocal microscopy.

The pores and membranes colocalized within the first 10 minutes. TAMRA then entered a much larger fraction of GUVs mixed with pores than GUVs without pores in control experiments (Fig. 3 a-c, g), demonstrating that the designed pores could facilitate transport across lipid membranes.

**Figure 3.**
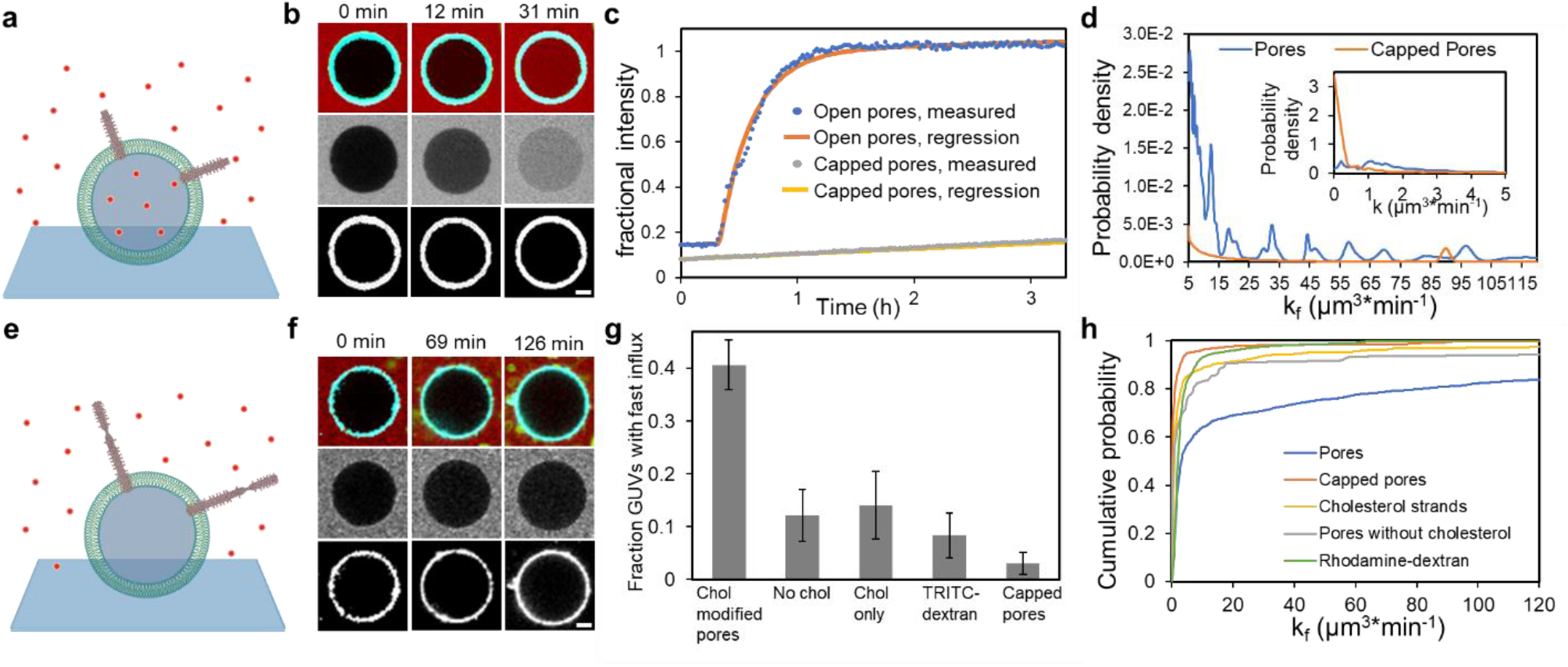
TAMRA transport through open and capped DNA origami pores. (**a**) Dye (TAMRA) influx assay through DNA pores into GUVs. (**b**) Fluorescence micrographs of an example GUV during TAMRA influx as in a. Top row: composite images of TAMRA (red), membrane (green), and DNA pores (cyan). Middle row: TAMRA fluorescence. Bottom row: DNA pore fluorescence. (**c**) Representative traces of fractional fluorescence intensities of GUVs during experiments as in and e. Pore-mediated influx begins after a lag period. Capped pores do not mediate influx. Regression curves of fractional intensities are best fits to equation (1) (Supplementary Note 10). (**d**) Distributions of GUV influx rates with DNA pores (N = 418) and capped DNA pores (N = 264). **e**. Dye influx assay through capped DNA pores into GUVs. (**f**) Fluorescence micrographs of an example GUV. Little TAMRA enters the vesicle. Fluorescent labeling as in (b) except DNA caps (green) are also in the top-row micrographs. (**g**) Fractions of GUVs with fast influx (*k*_*p*_>5 μm^3^/min) for (left to right) cholesterol modified pores (N=418), pores without cholesterol-DNA conjugates (N = 165), cholesterol-DNA conjugates (N = 114), pores and 500-kDa TRITC-dextran dye in place of TAMRA (N = 176) and capped pores. Error bars are 95% confidence intervals. (**h**) Cumulative distributions of influx rates into GUVs from the pore, capped pore, and control experiments in (g).

To quantify influx rates, we wrote computer programs to automatically identify 418 GUVs in micrographs and measured their internal fluorescence intensities over time. We computed the fractional fluorescence intensity (FFI) inside each vesicle, which we defined as the ratio of the mean fluorescence intensity inside a vesicle to the mean fluorescence intensity outside of vesicles in each field of view. The FFI of many vesicles initially increased slowly but then suddenly increased dramatically. Initial rates of FFI increase were similar to rates the FFI increases in control experiments (Fig. 3c), suggesting these initial increases were due to TAMRA slowly permeating through vesicle membranes. Although pores adsorbed to membranes rapidly, this initial phase often lasted over 30 minutes, suggesting slow pore insertion kinetics, consistent with the high energy barrier to DNA origami pore into GUVs observed previously.^36^

Just one event of sudden increase in FFI was observed for most GUVs, indicating either insertions of single pores or simultaneous insertion of multiple pores was most common. Multiple simultaneous insertions might occur if the insertion was a fusion event between a DNA pore-loaded SUV and a GUV. SUV byproducts remained after GUV preparation, and the low membrane curvatures of small vesicles can promote DNA origami pore insertions^36^. The pores were then transferred onto GUVs through vesicle fusions as vesicle fusions could frequently occur in solution, as has been observed previously^37,38^.

To better understand the rates of molecular transport, we developed a model that accounted for both fast, pore-mediated influx and slow influx due to TAMRA permeation across membranes. Fast TAMRA influx was modeled as net diffusion of TAMRA from the bulk solution outside the vesicle into a finite compartment through DNA pores (Supplementary Note 5). A DNA pore was modeled as a rigid cylindrical channel with non-permeable walls. Fick’s law predicted that the flux of TAMRA at a given time should be linearly proportional to the difference in concentrations inside and outside the pore and to the total cross-section area of the inserted channels.^34^ Slow, non-pore mediated TAMRA influx was modeled as a linear increase in fractional intensity (Supplementary Note 5):

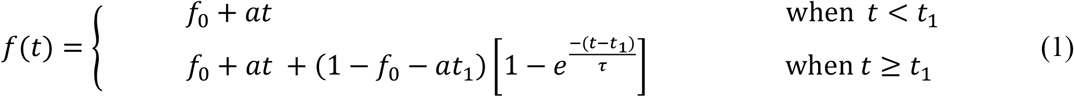

where *f*(*t*) is FFI at time *t. f*_0_ (initial FFI), *a, τ*, and *t*_1_ (time when fast influx starts) are constant parameters.

Non-linear regression produced good fits of the FFIs of most vesicles to equation (1); the small fraction of GUVs where the FFI increased in bursts, possibly due to lipid membrane defects^39^ were excluded. We used these fits to calculate the fast (pore-mediated) influx rate 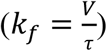, where *V* is vesicle volume (Supplementary Note 9), and the background influx rate (*k*_*b*_ = *aV*) for each vesicle.

*k*_*f*_ represents the total influx rate due to multiple inserted pores on each vesicle. Because DNA origami pores each have the same lengths and diameters and thus the same expected rate of pore-mediated transport, influx rates from different numbers of inserted pores should be quantized rather than continuously distributed, with each quantum being the rate of influx through a single pore. Thus, analysis of the distribution of measured *k*_*f*_ can reveal the single-pore transport rate.

We summed the posterior distributions of *k*_*f*_ (to include measurement error) for the GUVs to produce a composite probability distribution (Fig. 3d, Supplementary Note 10). The resulting distribution had a series of peaks. The Fourier transform of this distribution over 15 – 120 μm^3^/min had a dominant period of 13.1 ± 1.5 μm^3^/min (Supplementary Fig. 16). This measured value is very close to the first peak in the distribution (12.4 ± 2.5 μm^3^/min). Together these results suggest that each pore mediated TAMRA influx at 12.4 - 13.1 μm^3^/min, which corresponds to a net flux through each pore of 40.5 ± 4.6 molecules per second for the 309 nM TAMRA used in experiments (supplementary Note 6).

The expected rate of TAMRA influx through a single pore following Fickian diffusion is 14.7 ± 1.8 μm^3^/min (Supplementary Note 11), almost precisely the measured rate of 12.4 - 13.1 μm^3^/min. Although electrostatic interactions, van der Waals forces, and entry effects can play important roles in molecular transport in the nanoscale,^40^ the consistency between the measured TAMRA transport rates and the rates predicted by the continuum diffusion model suggested that the diffusion of TAMRA, an small uncharged molecule, in a DNA pore resembles molecular diffusion in a fluid continuum.

In the absence of pores, regression fits with equation (1) were almost uniformly below *k*_*f*_ of 5 μm^3^/min. In experiments with cholesterol-modified pores, a fraction of 0.41 ± 0.05 GUVs had *k*_*f*_>5 μm^3^/min (Fig. 3g), suggesting that cholesterol modification was necessary to induce fast influx. When TAMRA was replaced with the same concentration of a 500-kDa dye, tetramethylrhodamine isothiocyanate-dextran (TRITC-dextran) that has a hydrodynamic radius of 15.9 nm,^41^ a much lower percentage of GUVs (15%) had influx, demonstrating the size selectivity of the DNA pores.

Because small gaps exist between DNA helices in DNA origami structures,^42^ pores could mediate transport by allowing TAMRA to diffuse from one end of the pore to the other or by allowing TAMRA to diffuse through gaps in the pore channels’ helical walls. To determine whether TAMRA diffuses across channel walls, we measured the amount of flux mediated by pores bound to DNA origami caps (added in 2x excess). Fluorescence micrographs revealed that a fraction of 0.97 ± 0.01 of pores was uncharged bound to caps (Supplementary Figure 11). Capped pores localized to vesicle membranes, but fast TAMRA influx (>5 µm^3^/min) occurred in only a fraction of 0.03 ± 0.02 of GUVs (Fig. 3d-h). TAMRA therefore moves through pores via end-to-end diffusion and not through pore walls.

### Molecules transport end-to-end through longer DNA nanochannels

To investigate whether TAMRA could also diffuse across a much longer nanochannel without leaking across its walls, we grew DNA tile nanotubes from pores as nanochannels. 87% ± 3% of pores had nanochannels on them (Supplementary Fig. 6); their median length was 0.71 μm (95% CI = [0.64, 0.76] μm) (Fig. 4c). There were sharp increases of FFI in many of the 167 tracked vesicles after initial lag periods (Fig. 4a-b) in the dye influx experiment, implying that, like the DNA pores, DNA nanochannels mediated TAMRA flux across vesicle membranes.

**Figure 4.**
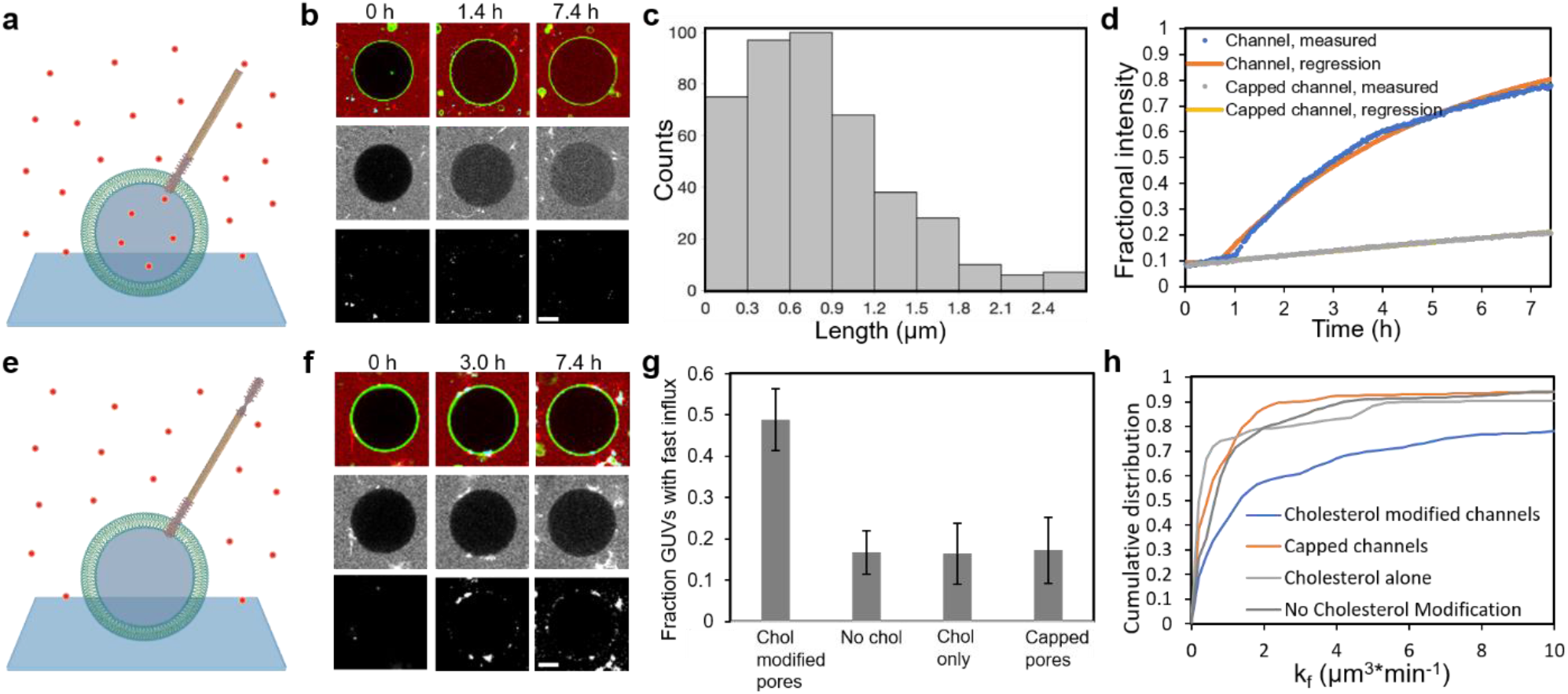
Rates of dye diffusion through DNA nanotube channels suggest end-to-end diffusion through nanochannels. (**a**) TAMRA dye influx assay through DNA nanochannels. (**b)** Fluorescence micrographs of an example GUV at different time points during TAMRA influx into the vesicle. Fluorescence labeling is as described in Fig. 3b, except nanochannels (red) are in the same fluorescence channel as TAMRA. The vesicle size increases, potentially from small osmotic pressure imbalances, were accounted for when calculating influx rates. (**c**) Histogram of nanochannel lengths. The last bin accounts for lengths >2.4 µm. (**d)** Representative fractional fluorescence intensity (FFI) traces of a GUV in the experiment in (a) and a GUV from the experiment in (e). Regression curves are best fits to a Fickian diffusion model (Supplementary Note 10). (**e**) TAMRA dye influx assay through capped nanochannels. (**f**) Fluorescence micrographs of an example GUV in the capped nanochannel experiment with the same fluorescence labeling as in (b) except DNA caps (green). (**g**) Fractions of GUVs with fast TAMRA influx for the experiments in (a), (e) and in control experiments where channels lack cholesterol modifications and with only cholesterol-DNA conjugates present. (**h**). Cumulative probability distribution of *k*_*f*_ between 0 – 10 μm^3^/min measured in the experiments in (g). Scale bars (b/f), 2 µm.

Diffusive flux through channels should be inversely proportional to channel length (Supplementary Note 7), so the flux through nanochannels should be smaller than through pores. To test this hypothesis, we fit the FFIs to equation (1). To collect as many measurements as possible, we included GUVs with two sequential insertion events in their FFI curves (Fig. 4d).

A fraction of 0.49 ± 0.08 vesicles had fast influx, *i.e*., *k*_*f*_>1 μm^3^/min, much higher than the fraction in control experiments with nanotubes without cholesterol modification (Fig. 4g). The cumulative distribution of channel-mediated influx rates (Fig. 4h) was also shifted higher for nanochannels than for controls, confirming that most vesicle influx was induced by inserted nanochannels.

Because the lengths of the nanochannels are polydisperse, they should have different influx rates. Consistent with this, the cumulative distribution of flux rates was not quantized. To compare the measured influx rates with those predicted by a bulk diffusive model, we compared the medians of each distribution. The measured median *k*_*f*_ for GUVs with fast influx in the dye influx experiment was 3.3 μm^3^/min (95% CI = [1.9, 5.7] μm^3^/min). This measured rate was somewhat larger than the theoretical medium-length channel influx rate of 1.3 μm^3^/min, possibly because this comparison does not account for the presence of multiple channels on a single vesicle.

The lower rate of TAMRA flux through nanochannels than through pores suggested that the TAMRA was diffusing primarily across channel lengths, rather than across channel walls. To determine the extent to which TAMRA diffused from one end of a channel to the other, we measured the extent to which capped channels could mediate TAMRA influx. A fraction of 0.90 ± 0.04 nanochannels were bound to caps after incubating the nanochannels with caps (Supplementary Fig. 17). Fewer than 20% of vesicles had levels of TAMRA influx consistent with channel-mediated transport (*k*_*f*_>1 μm^3^/min) in influx experiments with capped channels, indistinguishable from the percentage in control experiments (Fig. 4g). The cumulative distributions of *k*_*f*_ were also very close (Fig. 4h). DNA channel caps thus essentially eliminated most TAMRA flux through nanochannels, implying that the dominant mode of TAMRA flux was from one channel end to the other with minimum loss across channel walls.

### Evaluation of DNA-solute interactions through simulations

We used a combination of coarse-grained MD^31^ and Brownian dynamics simulations^43^ to probe microscopic parameters that govern diffusion of small molecules through the ends and the walls of the DNA origami pore. In our coarse-grained simulations, each bead represented at least four DNA base pairs. To mimic partial incorporation of the nanostructures in a lipid bilayer membrane, the second to last turns of all DNA helices were subject to an external attractive potential (Supplementary Note 12). Over the course of a multi-resolution coarse-grained simulation, the DNA pore was observed to retain, on average, its cylindrical shape and undergo local structural fluctuations, transiently opening slit-like passages through the walls of the pore located between DNA helices away from the interhelical crossovers (Supplementary Movie 4).

To evaluate conditions that facilitated diffusive end-to-end transport of small molecules, we transformed a representative equilibrated conformation of DNA pore into a 3D potential describing the interaction of a point-like particle (e.g., a dye molecule) with the DNA structure and a 3D diffusivity map prescribing the local diffusion constant of the particle (Supplementary Note 12). The continuum model of the DNA nanopore was combined with a planar potential representation of a lipid membrane to produce a model of the DNA pore embedded in a lipid bilayer (Fig. 5a). The model was surrounded by 396 point particles (corresponding to 4 mM concentration) and their diffusion in, around, and through the pore was simulated using the Brownian dynamics method.^44^

**Figure 5:**
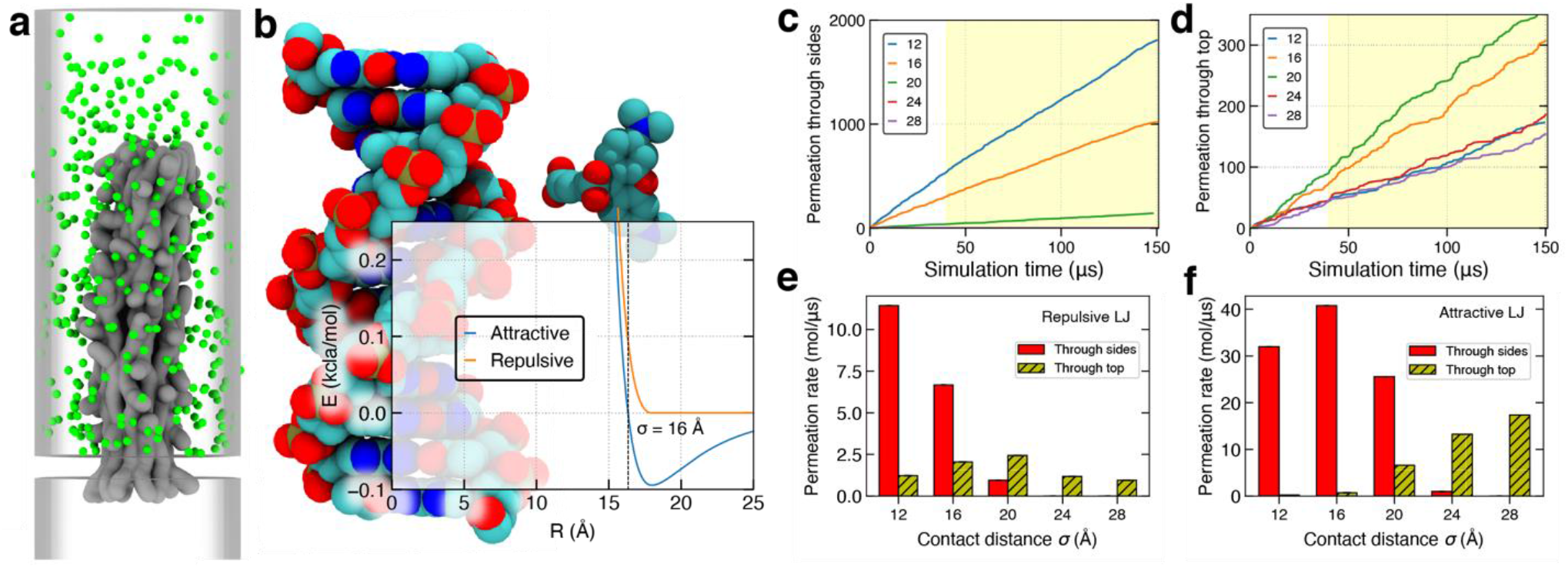
Brownian dynamics simulation of a small molecule permeation through the nanopore. **(a)** Representative configuration from a BD simulation of dye (green) diffusion through the nanopore structure. In the BD simulation of dye permeation, the bead-based representation of the nanostructure is replaced by grid-based interaction potential (gray; 1 kcal mol^−1^ isosurface) and a local diffusivity map. **(b)** Examples the attractive and repulsive interaction potential between DNA and a dye molecule. All-atom models of a DNA duplex and of a TAMRA dye are shown approximately to-scale in the background. Parameter σ defines effective contact distance. **(c, d)** The number of dye molecules permeated through the side walls (c) and through the top entrance (d) of the nanopore as a function of simulation time for purely repulsive models of dye–nanostructure interaction. The models vary by the value of the contact distance parameter σ. **(e)** The average rate of dye permeation computed using a linear regression fit to the data shown in panels c and d (shaded regions). The rate is much higher than in the experiment because of a much higher dye concentration used in the BD simulations. **(f)** The average rate of dye permeation for several models of attractive dye-DNA interaction.

By varying parameters describing the DNA-particle interaction (Fig. 5b), we determined effective interactions that could account for experimental observations. In a typical simulation, we counted the number of times a dye molecule entered the inner pore volume through the walls and through the top end of the pore. Fig. 5c and d show how the number of both types of permeation events changes over the course of the simulations. A linear regression fit to the plots yielded the permeation rate. For a purely repulsive potential, the permeation of dye molecules through the walls dominates over the end-to-end transport at effective contact distance, σ, of 16 Å or less (Fig. 5e). For σ > 20 Å, the rate of dye permeation through the end exceeds that through the walls. For σ = 24 and 28 Å, no through-the-wall permeation was observed, whereas molecules could still enter the nanopore through the end. Interestingly, adding attraction to DNA-dye interaction makes both permeation events more likely (Fig. 5f) and shifts the crossover from wall-to end-dominated transport to higher values of effective contact distance. Taken together, our modeling results indicate that end-to-end transport through the pore becomes dominant when, at contact, the distance between the center of the solute and the center of the DNA helix exceeds 2 nm.

## Discussion

In summary, we designed DNA origami pores and microns-long DNA nanochannels for transporting small molecules at rates predicted by the continuum diffusion model. The transport was shown to be end-to-end with minimal loss across channel walls and could be stopped by binding DNA caps to the channel’s end.

The self-assembled nanochannels studied here can self-repair^45^ and form interconnects between molecular landmarks^46^, suggesting how they might be used to form nanoscale flow networks for building artificial multicellular systems, widely used in studying cell-to-cell signaling. Large channel diameters could allow transport of macromolecules such as proteins and DNA. Long channel lengths would also be advantageous at characterizing the DNA-protein or protein-protein interactions within the channel for the long residence time of protein molecules transported through the channel. After functionalizing the channel interiors with antibodies or DNA aptamers, the nanotube channels can also serve as potential tools for detecting proteins of interest for biosensing applications.

## Materials and Methods

### Materials

All DNA oligodeoxynucleotides were purchased from Integrated DNA Technologies (IDT, Coralville, Iowa, USA), except the 7,240bp M13mp18 scaffold strand purchased from Bayou Biolabs (Los Angeles, CA, USA). Tris/acetate/EDTA (TAE) buffer was purchased from ThermoFisher Scientific (Waltham, MA, USA). 96-well glass-bottom plates with a streptavidin coating were purchased from Arrayit (Sunnyvale, CA, USA. Cat. No. M96S). 5(6)-Carboxytetramethylrhodamine (TAMRA) was purchased from Sigma-Aldrich (St. Louis, MO, USA). All lipids were purchased from Avanti Polar Lipids (Alabaster, AL, USA). Tetramethylrhodamine isothiocyanate–Dextran (TRITC-Dextran) of average molecular weight of 4,400 Da was purchased from Millipore Sigma (St. Louis, MO, USA. Cat. No. T1037).

### DNA pore assembly

A mixture containing 5 nM scaffold strand, 200 nM staple strand mix, and 25 nM attachment strand mix (Supplementary Note 1), was first prepared in TAE Mg^2+^ buffer (40 mM Tris-Acetate, 1 mM EDTA) with 12.5 mM magnesium acetate added (TAEM) (Supplementary Note 4.1). The assembly mixture was subjected to a thermal annealing ramp with an Eppendorf Mastercycler (Supplementary Note 3.3). The assembled pores were then purified to remove excessive DNA strands using a 100k MWCO Amicon Ultra centrifugal filter device (Millipore). Following purification, 0.15 μl of 100μM ATTO647-modified DNA strand that hybridizes to binding sites on attachment strands was added to 40 μl purified pore sample and incubated at room temperature for 15 min to allow fluorescently label the pores. The concentration of fluorescently labeled pores was approximately 1 nM, determined by measuring the concentration of a stock solution by counting the number of pores per field of view (86 μm × 86 μm) adsorbed to a glass slide from a specific reaction volume in fluorescence micrographs captured using a fluorescence microscope (Olympus IX71) with a 60×/1.45 NA oil immersion objective lens and a 1.6x magnifier lens. To modify the DNA pores with cholesterol, 1 μl DNA-cholesterol conjugate (Supplementary Note 3.6) at 10 μM was incubated at 50 °C for 10 minutes to alleviate cholesterol aggregation before being added to 40 μl purified DNA pores. The solution was then incubated for 10 minutes at room temperature.

### DNA nanotube channel assembly

DNA nanotube channels were self-assembled from DNA tiles and DNA pores. DNA tile monomers in an inactive form and DNA pores with adapters that presented monomer sticky ends were separately prepared before being mixed to form nanotube channels. Inactive DNA monomers were prepared by mixing six strands that fold into inactive monomers (Supplementary Note 2), each at 400 nM in TAEM and subjecting the mixture to a thermal annealing ramp (Supplementary Note 3.4). To make DNA pores with adapters, the nanopore assembly mixture was prepared as in the “DNA pore assembly” section except that 100 nM adapter strands were added to the mixture before thermal annealing (Supplementary Note 3.2). After assembly, the pores were purified and concentrated to 2 nM, then fluorescently labeled (Supplementary Note 3.4). 20 μl of pores were then mixed with 20 μl inactive monomers and 0.2 μl monomer activation strand (50 μM). The sample was then incubated at 37 °C for at least 15 hours to allow the nanotubes to grow. DNA-cholesterol conjugates were then hybridized to the pore regions of the nanotube channels by following the same steps used for modifying pores before influx assays.

### SUV Preparation

5 mg 1,2-diphytanoyl-sn-glycero-3-phosphocholine (DPhPC, Avanti Polar Lipids, USA) was dissolved in 1 ml chloroform in a glass vial. The solvent was evaporated by blowing nitrogen gas into the glass vial for 20 minutes so that a dry lipid film was formed. The glass vial was kept in a vacuum desiccator overnight to remove any residual solvent. 1 ml of 150 mM potassium chloride solution was then added to the glass vial to hydrate the lipid film. SUVs were then formed by 2-hour sonication in a bath sonicator (Bransonic Ultrasonic Cleaner (1510R-DTH)) at 40 °C. The SUVs were diluted 1,000 times with 150 mM potassium chloride before use.

### Transmission electron microscopy (TEM)

A formvar/carbon film support grid (Electron Microscopy Sciences, Hatfield, PA, USA, Cat No: FCF400-Cu) was glow discharged in a vacuum for 40 seconds. 10 µl of sample to be imaged was prepared (Supplementary Note 4). The grid was put onto the sample droplet and was incubated at room temperature for 5-10 minutes. The sample solution was then washed off by tapping the grid on MilliQ water droplets 5 times. The sample attached to the grid was stained by immediately tapping the grid on 2% w/w uranyl acetate (Electron Microscopy Sciences, Hatfield, PA, USA, Cat. No. 22400) solution droplets twice, followed by a 5-minute incubation on the uranyl acetate solution. The solution was then blotted, and the grid was first dried using a filter paper and then dried in air for at least an hour. The grid was imaged using a Field emission FEI Technai-12 electron microscope operated at 100 keV voltage.

### GUV Preparation

The protocol for preparing GUV was adapted from Horger et al.^47^ with modifications. 1% (w/w) agarose in deionized water was boiled in a microwave oven. 200 µl of the warm agarose solution was poured onto a glass-bottom dish. The dish was kept on a hotplate at 80 °C for an hour to form a dry agarose film. 1,2-diphytanoyl-sn-glycero-3-phosphocholine (DPhPC) lipids and 1,2-dioleoyl-sn-glycero-3-phosphoethanolamine-N-(biotinyl) (Biotinyl PE) were dissolved in chloroform to a concentration of 5 mg/ml in glass vials. A fluorescent lipid, 1-palmitoyl-2-(dipyrrometheneboron difluoride)undecanoyl-sn-glycero-3-phosphocholine (TopFluor® PC), was dissolved in chloroform to a concentration of 1 mg/ml in a glass vial. The three lipid solutions were mixed in chloroform to prepare a 1 mg/ml lipid mix solution with a composition of 88% DPhPC, 10% Biotinly PE, and 2% TopFluor PC. 40 µl lipid mix solution was then deposited onto the dry agarose film on the dish. The dish was kept in a vacuum desiccator overnight to allow the lipid solution to completely dry. After solvent was removed, 300 µl of 0.2 M sucrose solution was added to the film in the dish. The dish was kept undisturbed at room temperature for 2 days to allow GUV formation.

### Dye influx assay setup

90 µl TAE buffer supplemented with 0.2 M sucrose and 309 nM TAMRA was added into a well on a 96-well glass-bottom plate with a streptavidin coating. 6 µl GUV solution was then added into the well. The GUVs encapsulating 0.2 M sucrose settled down onto and became immobilized on the glass surface in about 5 minutes due to density differences between the solutions inside and outside the GUVs and biotin-streptavidin binding between the vesicles and the glass surface. 35 µl of solution containing either pores, capped pores, nanotube channels, or capped nanotube channels prepared in TAEM buffer, after cholesterol modification, were mixed with 9 µl 1 M glucose before being added into the well to maintain a final concentration of 0.2 M glucose. The final concentration of magnesium ions was 3.1 mM.

The sample on the glass-bottom plate was imaged on an inverted confocal microscope (Nikon Ti2-E with A1 confocal unit) using a 60x/1.49 NA oil immersion objective lens. DNA pores were imaged using a 640 nm diode laser, TAMRA and DNA nanotubes were imaged using a 560 nm diode laser, and GUVs were imaged using a 480 nm diode laser. All three fluorescent channels at one focal plane position in a large area of 4 × 4 fields of view were captured with 60- or 71-second time intervals. Each field of view had dimensions of 512 × 512 pixels, corresponding to a physical field-of-view size of 210 µm × 210 µm. The experiments with DNA pores were observed for a total time length of 210 minutes and experiments with DNA nanotubes were observed for a total time length of 8 hours. The Nikon Perfect Focus System (PFS) was used to eliminate axial focus fluctuations.

### Fluorescence image data processing

To quantify the fluorescence intensities of TAMRA over time inside and outside GUVs observed using confocal microscopy, we first used ImageJ software to identify and track individual GUVs captured. The outlines of GUVs at each time point were identified by applying the “thresholding” function in ImageJ software to convert the grayscale images of GUVs into binary images. The image areas inside GUVs at each time point were identified by selecting the areas separated by GUV outlines that had circularities greater than 0.65 using the “Analyze Particles” function in ImageJ software. Similarly, one area outside GUVs at each time point was identified by selecting the largest area in the image with circularity smaller than 0.6. The mean fluorescence intensities of TAMRA in the areas inside and outside the identified GUVs at each time point were measured from grayscale images of TAMRA. The TAMRA concentration fractions of individual GUVs at each time point were calculated by dividing the mean intensities inside GUVs to the mean intensity outside GUVs.

## Acknowledgment

Y.L. and R.S. acknowledge support from the Defense Advanced Research Projects Agency (D16AP00147) and the Department of Energy (DE-SC0010426). C.M., H.J. and A.A. acknowledge support from the National Science Foundation of USA (DMR-1827346) and the supercomputer time provided through the XSEDE allocation grant (MCA05S028) and the Leadership Resource Allocation MCB20012 on Frontera of the Texas Advanced Computing Center. We sincerely thank Sisi Jia, Naresh Niranjan, and Samuel Schaffter for useful discussions.

## Author Contributions

Y.L. and R.S. planned and designed the experiments. Y.L. conducted the experiments. Y.L. and B.M. performed data analysis. C.M., H.J., and A.A. performed simulations. Y.L., R.S., C.M., H.J., and A.A. wrote the paper. All authors discussed the results and commented on the manuscript. H.J. was affiliated with Department of Physics, University of Illinois Urbana-Champaign, Urbana, IL at the time of conducting this study and is currently affiliated with Department of Biotechnology, Indian Institute of Technology Hyderabad, Kandi, Sangareddy, Telangana, India

## Competing financial interests

The authors declare no competing financial interests.

## Data and materials availability

All data needed to evaluate the conclusions in the paper are present in the paper and/or the Supplementary Materials.

## Additional information

**Supplementary Information** accompanies this paper includes:

Supplementary Information (Supplementary Note 1-11, Supplementary Table 1-4, Supplementary Figure 1-18, and Supplementary References)

Supplementary Movies 1-5

## Notes

### Competing Interest Statement

The authors have declared no competing interest.

## References

1. Kowalczyk, S. W., Blosser, T. R. & Dekker, C. Biomimetic nanopores: Learning from and about nature. Trends in Biotechnology vol. 29 607–614 (2011).

2. de Riccardis, F., Izzo, I., Montesarchio, D. & Tecilla, P. Ion transport through lipid bilayers by synthetic ionophores: Modulation of activity and selectivity. Accounts of Chemical Research 46, 2781–2790 (2013).

3. Hou, X. et al. A Biomimetic Potassium Responsive Nanochannel: G-Quadruplex DNA Conformational Switching in a Synthetic Nanopore. Journal of the American Chemical Society 131, 7800–7805 (2009).

4. Maingi, V., Lelimousin, M., Howorka, S. & P Sansom, M. S. Gating-like Motions and Wall Porosity in a DNA Nanopore Scaffold Revealed by Molecular Simulations. (2015) doi:10.1021/acsnano.5b06357.

5. Zhao, K. & Wu, H. Structure-dependent water transport across nanopores of carbon nanotubes: Toward selective gating upon temperature regulation. Physical Chemistry Chemical Physics 17, 10343–10347 (2015).

6. Liu, L. & Wu, H.-C. DNA-Based Nanopore Sensing. Angewandte Chemie International Edition 55, 15216–15222 (2016).

7. Bayley, H. & Cremer, P. S. Stochastic sensors inspired by biology. Nature vol. 413 226– 230 (2001).

8. Shi, W., Friedman, A. K. & Baker, L. A. Nanopore Sensing. Analytical Chemistry vol. 89 157–188 (2017).

9. Koçer, A., Walko, M., Meijberg, W. & Feringa, B. L. Chemistry: A light-actuated nanovalve derived from a channel protein. Science 309, 755–758 (2005).

10. Duan, R., Xia, F. & Jiang, L. Constructing tunable nanopores and their application in drug delivery. ACS Nano vol. 7 8344–8349 (2013).

11. Gokel, G. W. & Negin, S. Synthetic ion channels: From pores to biological applications. Accounts of Chemical Research 46, 2824–2833 (2013).

12. Montenegro, J., Ghadiri, M. R. & Granja, J. R. Ion channel models based on self-assembling cyclic peptide nanotubes. Accounts of Chemical Research 46, 2955–2965 (2013).

13. Neevel, J. G. & Nolte, R. J. M. Ion transport across vesicle bilayers mediated by an artificial channel compound. Tetrahedron Letters 25, 2263–2266 (1984).

14. Langecker, M. et al. Synthetic lipid membrane channels formed by designed DNA nanostructures. Science 338, 932–936 (2012).

15. Kawano, R. et al. Metal-Organic Cuboctahedra for Synthetic Ion Channels with Multiple Conductance States. Chem 2, 393–403 (2017).

16. Fyles, T. M. & Tong, C. C. Long-lived and highly conducting ion channels formed by lipophilic ethylenediamine palladium(II) complexesw. (2006) doi:10.1039/b610660a.

17. Liu, B., Li, X., Li, B., Xu, B. & Zhao, Y. Carbon nanotube based artificial water channel protein: Membrane perturbation and water transportation. Nano Letters 9, 1386–1394 (2009).

18. Krishnan, S. et al. Molecular transport through large-diameter DNA nanopores. Nature Communications 7, 1–7 (2016).

19. Göpfrich, K. et al. Large-Conductance Transmembrane Porin Made from DNA Origami. ACS Nano 10, 8207–8214 (2016).

20. Thomsen, R. P. et al. A large size-selective DNA nanopore with sensing applications. Nature Communications 10, 1–10 (2019).

21. Diederichs, T. et al. Synthetic protein-conductive membrane nanopores built with DNA. Nature Communications 10, 1–11 (2019).

22. Iwabuchi, S., Kawamata, I., Murata, S. & Nomura, S. I. M. A large, square-shaped, DNA origami nanopore with sealing function on a giant vesicle membrane. Chemical Communications 57, 2990–2993 (2021).

23. Yang, Q., Guo, Z., Liu, H., Peng, R., Xu, L., Bi, C., He, Y., Liu, Q., &amp; Tan, W. A cascade signaling network between artificial cells switching activity of synthetic transmembrane channels. Journal of the American Chemical Society, 143(1), 232–240. (2020).

24. Barati Farimani, A., Dibaeinia, P., &amp; Aluru, N.R. DNA origami–graphene hybrid nanopore for DNA detection. ACS Applied Materials &amp; Interfaces, 9(1), 92–100. (2016).

25. Shen, B., Piskunen, P., Nummelin, S., Liu, Q., Kostiainen, M. A., &amp; Linko, V. Advanced DNA nanopore technologies. ACS Applied Bio Materials, 3(9), 5606–5619. (2020).

26. Iwabuchi, S., Kawamata, I., Murata, S., &amp; Nomura S.-ichiro M. A large, square-shaped, DNA origami nanopore with sealing function on a giant vesicle membrane. Chemical Communications, 57(24), 2990–2993. (2021).

27. Sharma, B., Moghimianavval, H., Hwang, S.-W., &amp; Liu, A.P. Synthetic cell as a platform for understanding membrane-membrane interactions. Membranes, 11(12), 912. (2021).

28. Howorka, S. Building membrane nanopores. Nature Nanotechnology vol. 12 619–630 (2017).

29. Zhao, Z., Yan, H., &amp; Liu, Y. A route to scale up DNA origami using DNA tiles as folding staples. Angewandte Chemie, 122(8), 1456–1459 (2010).

30. Mohammed, A. M. & Schulman, R. Directing self-assembly of DNA nanotubes using programmable seeds. Nano Letters 13, 4006–4013 (2013).

31. Maffeo, C. & Aksimentiev, A. MrDNA: a multi-resolution model for predicting the structure and dynamics of DNA systems. Nucleic Acids Research 48, (2020).

32. Agrawal, D. K. et al. Terminating DNA Tile Assembly with Nanostructured Caps. ACS Nano 11, 9770–9779 (2017).

33. Continuum Theory of Diffusion. in 27–36 (Springer, Berlin, Heidelberg, 2007). doi:10.1007/978-3-540-71488-0_2.

34. Flynn, G. L., Yalkowsky, S. H. & Roseman, T. J. Mass transport phenomena and models: Theoretical concepts. Journal of Pharmaceutical Sciences vol. 63 479–510 (1974).

35. Wang, L. et al. A general strategy to develop cell permeable and fluorogenic probes for multicolour nanoscopy. Nature Chemistry 12, 165–172 (2020).

36. Burns, J. R. & Howorka, S. Defined Bilayer Interactions of DNA Nanopores Revealed with a Nuclease-Based Nanoprobe Strategy. ACS Nano 12, 3263–3271 (2018).

37. Haluska, C. K. et al. Time scale of membrane fusion revealed by direct imaging of vesicle fusion with high temporal resolution. Proceedings of the National Academy of Sciences of the United States of America 103, 15841–15846 (2006).

38. Chen, L., Liang, S., Chen, Y., Wu, M., Zhang, Y. (2019). Destructing the Plasma Membrane with Activatable Vesicular DNA Nanopores. ACS Applied Materials 12(1), 96–105. https://doi.org/10.1021/acsami.

39. Karatekin, E. et al. Cascades of transient pores in giant vesicles: Line tension and transport. Biophysical Journal 84, 1734–1749 (2003).

40. Sparreboom, W., van den Berg, A. & Eijkel, J. C. T. Principles and applications of nanofluidic transport. Nature Nanotechnology vol. 4 713–720 (2009).

41. Armstrong, J. K., Wenby, R. B., Meiselman, H. J. & Fisher, T. C. The hydrodynamic radii of macromolecules and their effect on red blood cell aggregation. Biophysical Journal 87, 4259–4270 (2004).

42. Fischer, S. et al. Shape and interhelical spacing of DNA origami nanostructures studied by small-angle X-ray scattering. Nano Letters 16, 4282–4287 (2016).

43. Comer, J. & Aksimentiev, A. Predicting the DNA Sequence Dependence of Nanopore Ion Current Using Atomic-Resolution Brownian Dynamics. The Journal of Physical Chemistry C 116, 3376–3393 (2012).

44. Ermak, D. L. & McCammon, J. A. Brownian dynamics with hydrodynamic interactions. The Journal of Chemical Physics 69, 1352–1360 (1978).

45. Li, Y. & Schulman, R. DNA Nanostructures that Self-Heal in Serum. Nano Letters 19, 3751–3760 (2019).

46. Mohammed, A. M., Šulc, P., Zenk, J. & Schulman, R. Self-assembling DNA nanotubes to connect molecular landmarks. (2016) doi:10.1038/NNANO.2016.277.

47. Horger, K. S., Estes, D. J., Capone, R. & Mayer, M. Films of agarose enable rapid formation of giant liposomes in solutions of physiologic ionic strength. Journal of the American Chemical Society 131, 1810–1819 (2009).

