## Supplementary Information for "Lossless end-to-end transport of small molecules through micron-length DNA nanochannels"

**This PDF file includes:**

Supplementary Note 1 to 12

Supplementary Figures S1 to S18

Supplementary Movies captions S1 to S5

References 1 to 13

**Other Supplementary Materials for this manuscript include the following:**

Supplementary Movies S1 to S5

### Table of contents

**Supplementary Note 1** – Design of the DNA nanopore

**Supplementary Figure 3** – Staple map for the DNA nanopore

**Supplementary Table 1** – Names for the 72 staple strands of the DNA nanopore

**Supplementary Note 2** – DNA nanotube design

**Supplementary Figure 2** –Schematic of the monomer activation reaction

**Supplementary Figure 3** – Design of the DNA cap

**Supplementary Table 2** – DNA cap staple sequences

**Supplementary Note 3** – DNA nanopore and nanotube channel preparation

**Supplementary Figure 4** – All-atom model of the DNA nanopore structure obtained through multi-resolution simulations

**Supplementary Figure 5** – A structural snapshot from a coarse-grained model of the DNA cap

**Supplementary Figure 6** – Example fluorescence micrographs of DNA nanotube channels attached to DNA pores

**Supplementary Note 4** – Preparation of samples for Transmission Electron Microscopy (TEM)

**Supplementary Figure 7** – Additional TEM Images of DNA origami nanopores without added DNA-cholesterol conjugates

**Supplementary Figure 8** – Additional TEM Images of DNA nanopores interacting with SUVs

**Supplementary Figure 9** – Additional TEM Images of DNA origami caps

**Supplementary Figure 10** – TEM Images of capped nanopores

**Supplementary Figure 11** – Example wide-field fluorescence image of DNA caps bound to DNA nanopores

**Supplementary Note 5** – A bulk diffusion model of TAMRA influx into vesicles through DNA nanopores

**Supplementary Note 6** – Rate of TAMRA influx through a single DNA nanopore

**Supplementary Note 7** – A diffusion model for transport of TAMRA through DNA nanotube channels in the influx assays

**Supplementary Note 8** – Image analysis of fluorescence micrographs from dye influx experiments

**Supplementary Figure 12** – Example fluorescence micrographs of GUVs from the dye influx experiment with DNA nanopores.

**Supplementary Note 9** – Calculating vesicle volumes in the dye influx experiments

**Supplementary Note 10** – Regression analysis of fractional intensity data

**Supplementary Figure 13** – Example plots of measured fractional intensities of GUVs and corresponding regression curves

**Supplementary Table 3** – Kinetic parameters for vesicles in Supplementary Figure 13

**Supplementary Figure 14** – Example plots of measured and fitted fractional intensities of GUVs in experiments where DNA nanochannels are added to GUVs

**Supplementary Table 4** – Kinetic parameters for vesicles in Supplementary Figure 14

**Supplementary Figure 15** – Measured density distributions of the influx rates in the nanochannel and capped nanochannel dye influx experiments

**Supplementary Note 11** – Theoretical rate of TAMRA transport through a single DNA pore

**Supplementary Figure 16** – Fourier transform of the probability density distribution of influx rates of TAMRA into DNA pores

**Supplementary Figure 17** – Example wide-field fluorescence image of DNA caps bound to DNA nanochannels

**Supplementary Note 12:** Computer simulation of small molecule diffusion through a DNA pore

**Supplementary Figure 18:** Interaction potential between two dye molecules used in the Brownian dynamics simulations of dye diffusion through the pore structure.

**Supplementary Movie 1.** DNA nanopores induce fast TAMRA influx into a GUV.

**Supplementary Movie 2.** DNA nanochannels attached to DNA pores insert onto GUV membranes.

**Supplementary Movie 3.** DNA nanochannels (red) induce TAMRA (red) influx into a GUV.

**Supplementary Movie 4:** Multi-resolution equilibration of DNA seed nanopore embedded in a lipid bilayer membrane.

**Supplementary Movie 5.** Brownian dynamics simulation of dye diffusion through the pore.

**Supplementary References**

#### Supplementary Note 1. Design of the DNA nanopore

The DNA nanopore design was adapted from the pore structure previously published as the nanotube seed structure by Mohammed & Schulman[1].

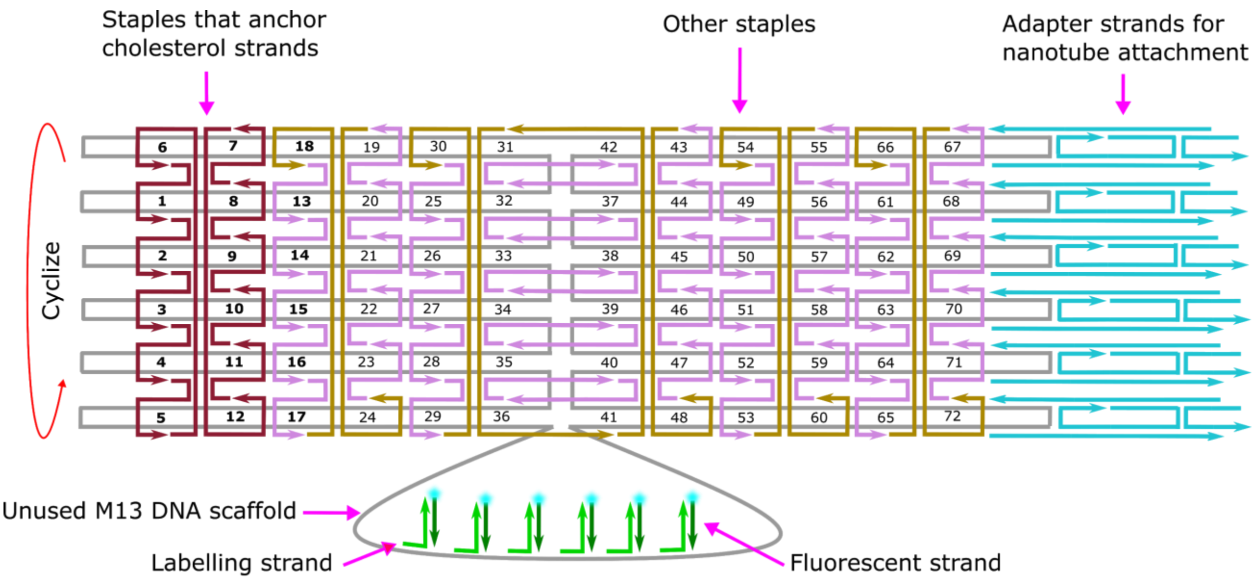


**Supplementary Figure 1. Staple map showing the positions of 72 staple strands, 24 adapter strands, and the unused M13 scaffold for the DNA origami nanopore.** The 12 staples on the left side of the map (dark red) were modified to present 15-nt overhangs that allow cholesterol-modified strands to bind. The other 60 staples (lavender and brown) have hairpin domains to help the origami to correctly cyclize during thermal annealing, as described in Mohammed & Schulman[1]. The overhangs for binding of cholesterol strands and hairpin domains are not shown on the strands in the map. To visualize the DNA origami using fluorescence microscopy, the DNA origami is labelled with fluorophores attached to fluorescent labeling strands that can bind to the unused region of M13 scaffold (gray)[1]. The seed adapter strands (cyan) present the sticky end sequences of the monomers used this study for nanotube growth on the DNA origami.

**Supplementary Table 1**. Names for the 72 staple strands of the DNA nanopore.

| | | | | | | | | | | | |
|---|---|---|---|---|---|---|---|---|---|---|---|
| **T_5R2F_CholDNA.Tag1** | **T_5R2E_CholDNA.Tag7** | **T_3R2F_HP** | **T_3R2E_HP** | **T_1R2F_HP** | **T_1R2E_HP** | **T1R2F_HP** | **T1R2E_HP** | **T3R2F_HP** | **T3R2E_HP** | **T5R2F_HP** | **T5R2E_HP** |
| **T_5R4F_CholDNA.Tag2** | **T_5R4E_CholDNA.Tag8** | **T_3R4F_HP** | **T_3R4E_HP** | **T_1R4F_HP** | **T_1R4E_HP** | **T1R4F_HP** | **T1R4E_HP** | **T3R4F_HP** | **T3R4E_HP** | **T5R4F_HP** | **T5R4E_HP** |
| **T_5R6F_CholDNA.Tag3** | **T_5R6E_CholDNA.Tag9** | **T_3R6F_HP** | **T_3R6E_HP** | **T_1R6F_HP** | **T_1R6E_HP** | **T1R6F_HP** | **T1R6E_HP** | **T3R6F_HP** | **T3R6E_HP** | **T5R6F_HP** | **T5R6E_HP** |
| **T_5R8F_CholDNA.Tag4** | **T_5R8E_CholDNA.Tag10** | **T_3R8F_HP** | **T_3R8E_HP** | **T_1R8F_HP** | **T_1R8E_HP** | **T1R8F_HP** | **T1R8E_HP** | **T3R8F_HP** | **T3R8E_HP** | **T5R8F_HP** | **T5R8E_HP** |
| **T_5R10F_CholDNA.Tag5** | **T_5R10E_CholDNA.Tag11** | **T_3R10F_HP** | **T_3R10E_HP** | **T_1R10F_HP** | **T_1R10E_HP** | **T1R10F_HP** | **T1R10E_HP** | **T3R10F_HP** | **T3R10E_HP** | **T5R10F_HP** | **T5R10E_HP** |
| **T_5R12F_CYC_CholDNA.Tag6** | **T_5R12E_CYC_CholDNA.Tag12** | **T_3R12F_HP** | **T_3R12E_CYC_HP** | **T_1R12_CYC_HP** | **T_1R12E_CYC_HP** | **T_1R12F_CYC_HP** | **T1R12E_CYC_HP** | **T3R12F_CYC_HP** | **T3R12E_CYC_HP** | **T5R12F_CYC_HP** | **T5R12E_CYC_HP** |

The sequences of the 12 staple strands to which cholesterol modified strands hybridize ("cholesterol_strand") are listed below.

| | |
|---|---|
| T_5R2F_CholDNA.Tag1 | TTGCAGCTACGTCTATGAGTTTCAAAGGAACGTCCACCGTTTTCGGTGGACTTA |
| T_5R4F_CholDNA.Tag2 | TTGCAGCTACGTCTAAAAAAAGGCTTTTGCGGTGGTCCGTTTTCGGACCACTTG |
| T_5R6F_CholDNA.Tag3 | TTGCAGCTACGTCTAACGGCTACAAGTACAACTCGGCACTTTTGTGCCGAGTTC |
| T_5R8F_CholDNA.Tag4 | TTGCAGCTACGTCTAGCTCCATGACGTAACACGGATCGCTTTTGCGATCCGTTA |
| T_5R10F_CholDNA.Tag5 | TTGCAGCTACGTCTAACGAGTAGATCAGTTGCACCGCTGTTTTCAGCGGTG TTA |
| T_5R12F_CYC_CholDNA.Tag6 | TTGCAGCTACGTCTAGGAATTACCACCACCCGTGAGGCGTTTTCGCCTCACTTT |
| T_5R2E_CholDNA.Tag7 | TTGCAGCTACGTCTAGAGAATAGGTCACCAGCGGAACCGTTTTCGGTTCCGTTT |
| T_5R4E_CholDNA.Tag8 | TTGCAGCTACGTCTAAAAGGCCGCTCCAAAACCGTGGCGTTTTCGCCACGGTTG |
| T_5R6E_CholDNA.Tag9 | TTGCAGCTACGTCTAGCGAAACAAGAGGCTTGTGCTGCGTTTTCGCAGCACTTT |
| T_5R8E_CholDNA.Tag10 | TTGCAGCTACGTCTACCAAATCATTACTTAGACGCTGGCTTTTGCCAGCGTTTC |
| T_5R10E_CholDNA.Tag11 | TTGCAGCTACGTCTAAAAGATTCTAAATTGGCGACGGACTTTTGTCCGTCGTTG |
| T_5R12E_CYC_CholDNA.Tag12 | TTGCAGCTACGTCTACTCAGAGCGAGGCATAGGCTCCGCTTTTGCGGAGCCTTG |

The sequences of the remaining 60 staple strands are those of the same names reported in Mohammed & Schulman[1]. The 24 adapter strands, added along with staple strands at annealing, were the same as the A adapters reported in Jia et al[2].

DNA nanopores were labeled with ATTO647 fluorophore dyes for fluorescence imaging. The labeling system consists of 100 attachment strands, each of which contains a subsequence that binds to the section of the M13mp18 scaffold that is not folded by staples. The remainder of each attachment strand binds to a labeling strand that has ATTO647 fluorophore dye on the 5' end, "labeling_strand_ATTO647N". The sequences of the attachment strands are the same as those listed in Mohammed & Schulman[1].

**cholesterol_strand**: TAGACGTAGCTGCAA/3CholTEG/
**labeling_strand_ATTO647N**: /5ATTO647NN/AAGCGTAGTCGGATCTC

/3CholTEG/ denotes a cholesterol molecule conjugated to the 3' end of the DNA strand. /5ATTO647NN/ denotes a ATTO647N (NHS ester) fluorophore molecule conjugated to the 5' end of the DNA strand.

#### Supplementary Note 2. DNA nanotube design

The DNA nanotubes are formed from the polymerization of oligomeric DNA monomers; nanotubes can either grow from or attach to DNA pores to form extended nanochannels with one end that can traverse a membrane. The monomer design and sequences in this study were adapted those used to assemble DNA nanotubes in a previous study[3]. To ensure that monomers do not start to assemble into nanotubes before they are mixed with DNA pores, we designed modified monomers that could be assembled in an inactive form during annealing and [5]could then be activated, i.e. reach a conformation that allowed assembly into nanotubes, by a strand-displacement reaction with an activation strand (Supplementary Figure 2). These monomers were based on those reported in Zhang et al[4].

One of the sticky ends of the inactive monomers is double-stranded, which prevents the monomers from forming a lattice by sticky end joining. The activation strand, 'SEs_activation', upon addition to the solution, displaces the 'SEs_inactive_strand5_right' strand and exposes a single-stranded sticky end where a double-stranded end was previously. The resulting products have four exposed sticky ends, allowing assembly of DNA nanotubes.

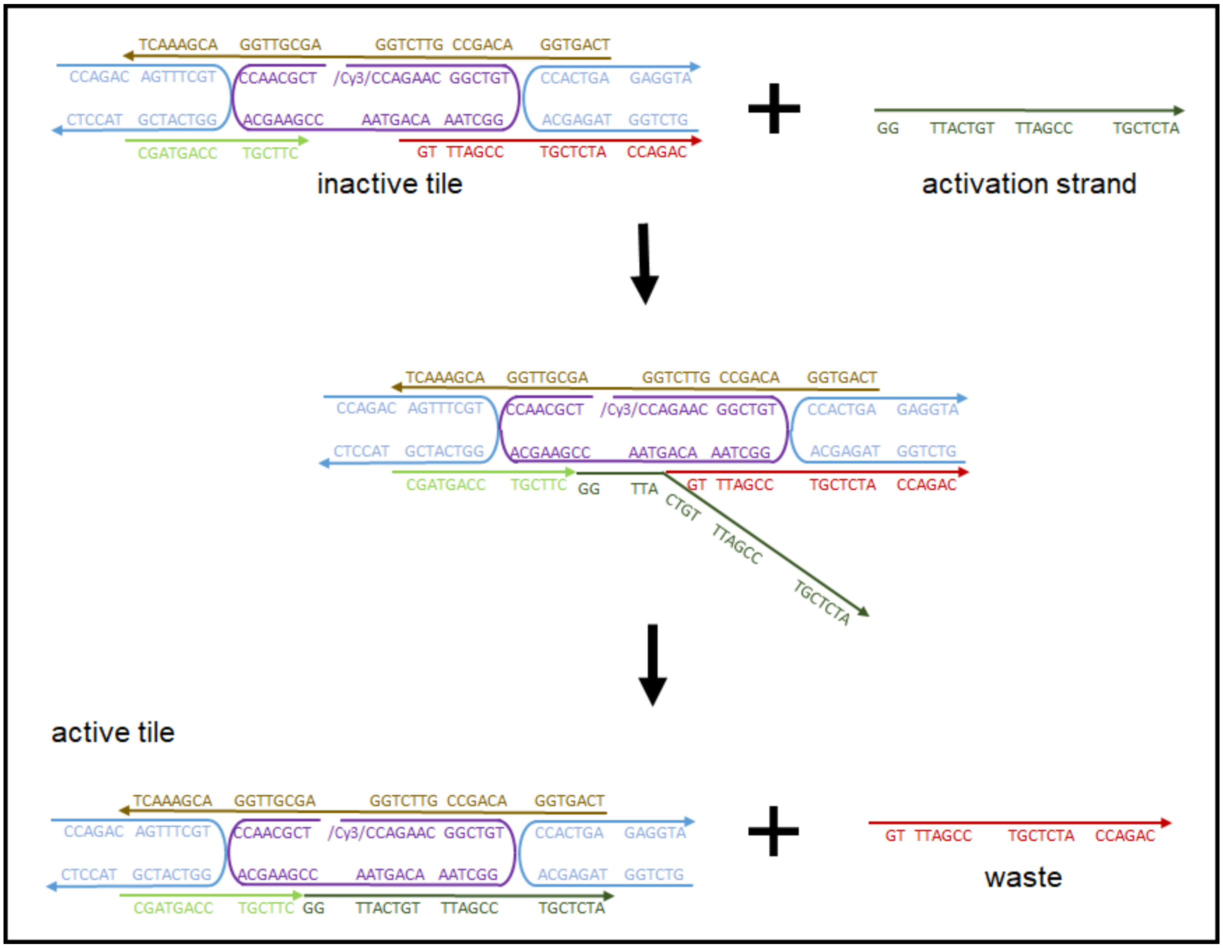


**Supplementary Figure 2. Schematic of the monomer activation reaction.** The activation strand displaces the strand that covers one of the sticky ends to active the inactive monomer.

DNA monomer sequences:

**SEs_1**: TCAGTGGACAGCCGTTCTGGAGCGTTGGACGAAACT
**SEs_2**: CCAGACAGTTTCGTGGTCATCGTACCTC
**SEs_3-5'Cy3**: /CY3/CCAGAACGGCTGTGGCTAAACAGTAACCGAAGCACCAACGCT
**SEs_4**: GTCTGGTAGAGCACCACTGAGAGGTA
**SEs_inactive_strand5_left**: CGATGACCTGCTTC
**SEs_inactive_strand5_right**: GTTACTGTTTAGCCTGCTCTACCAGAC
**SEs_activation**: GGTTACTGTTTAGCCTGCTCTA

/Cy3/ denotes a Cy3 fluorophore covalently attached to the 5' end of DNA.

#### Supplementary Table 2. DNA cap staple sequences

We designed the DNA cap by replacing 12 staple strands in the DNA origami structure in Mohammed & Schulman[1] while preserving the other 60 staple strands.

| **Name** | **Replaced staple name** | **Sequence** |
|---|---|---|
| stp_47_1 | T_5R12E_CYC_HP | CTCAGAGCGAGGCATAGGCTCCGCTTTTGCGGAGCCTTGTAAGAGCCCGCCACC |
| stp_47_2 | T_5R2E_HP | GAGAATAGGTCACCAGCGGAACCGTTTTCGGTTCCGTTTACAAACTACAGGTAG |
| stp_47_3 | T_5R4E_HP | AAAGGCCGCTCCAAAACCGTGGCGTTTTCGCCACGGTTGGAGCCTTTTCATTAC |
| stp_47_4 | T_5R6E_HP | GCGAAACAAGAGGCTTGTGCTGCGTTTTCGCAGCACTTTGAGGACTTACCAAGC |
| stp_47_5 | T_5R8E_HP | CCAAATCATTACTTAGACGCTGGCTTTTGCCAGCGTTTCCGGAACGAGGGAGTT |
| stp_47_6 | T_5R10E_HP | AAAGATTCTAAATTGGCGACGGACTTTTGTCCGTCGTTGCTTGAGAAGCGGAGT |
| stp_79_1 | T_3R12E_CYC_HP | CCCTCAGATCGTTTACCGCTTGCGTTTTCGCAAGCGTTCAGACGACTACCGCCA |
| stp_79_2 | T_3R2E_HP | TGCTAAACTCCACAGAGCCAGTGCTTTTGCACTGGCTTCAGCCCTCTTAATAAA |
| stp_79_3 | T_3R4E_HP | ATATATTCTCAGCTTGCCGTCCGCTTTTGCGGACGGTTCTTTCGAGTCATCAAG |
| stp_79_4 | T_3R6E_HP | CTCATCTTGGAAGTTTCGGATGGCTTTTGCCATCCGTTCCATTAAACTAAAACA |
| stp_79_5 | T_3R8E_HP | AGTAATCTTCATAAGGTCTGGTCGTTTTCGACCAGATTGAACCGAACATAACCG |
| stp_79_6 | T_3R10E_HP | ACGAACTATTAATCATGGCACCTGTTTTCAGGTGCCTTTGTGAATTTGGGATTT |

**Supplementary Note 3. DNA nanopore and nanotube channel preparation**

The protocols for assembling DNA nanopores and nanotube channels were adapted from the methods reported in Li & Schulman[3].

**3.1 Preparation of annealing solution for DNA nanopores**

To assemble DNA nanopores, 50 µl of an annealing mixture was prepared that contained M13mp18 scaffold, staple strands, and fluorescence attachment strands in TAEM buffer in the quantities shown below.

Recipe for preparing pore annealing solution:

| | Desired final concentration (nM or fold) | Stock concentration (nM or fold) | Volume added (µl) |
|---|---|---|---|
| $H_2O$ | | | 34 |
| TAEM buffer | 1 | 10 | 5 |
| Staple mix (concentrations are per strand) | 200 | 1389 | 7.2 |
| M13 scaffold | 5 | 100 | 2.5 |
| Fluorescence attachment strand mix (concentrations are per strand) | 25 | 1000 | 1.3 |
| Total | | | 50 |

**3.2 Preparation of annealing solution for DNA nanopores with adapters**

To allow DNA monomers to assemble on DNA nanopores to form nanotube channels, 24 adapter strands were added to the nanopore structures as they were assembled. The design and sequences of the adapter strands added to the nanopores are the Adapter A strands described in Jia et al[2].

50 μl annealing mixture consisting of M13mp18 scaffold, staple strands, adapter strands, and fluorescence attachment strands in TAEM buffer were prepared in the quantities shown below.

Recipe for preparing the annealing mix of nanopores for nanotube attachment:

| | Desired final concentration (nM or fold) | Stock concentration (nM or fold) | Volume added (μl) |
|---|---|---|---|
| $H_2O$ | | | 29 |
| 10x TAEM buffer | 1 | 10 | 5 |
| Nanopore staple mix (concentrations are per strand) | 200 | 1389 | 7.2 |
| Adapter A strand mix (concentrations are per strand) | 100 | 1000 | 5 |
| M13 scaffold | 5 | 100 | 2.5 |
| Fluorescent attachment strand mix (concentrations are per strand) | 25 | 1000 | 1.3 |
| Total | | | 50 |

**3.3 Preparation of annealing solution for DNA channel caps**

The composition of the solution annealed to form DNA channel caps is the same as the composition of the solution for annealed to form DNA nanopores described in Supplementary Note 3.2, except that DNA cap staple mix (Supplementary Table 2) was used instead of nanopore staple mix and Adapter B strand mix was used instead of Adapter A strand mix. Adapter B strands create a facet on the assembled structures presenting sticky ends that hybridize to the sticky ends on the facet formed by Adapter A strands. The designs and sequences for Adapter B strands are described in Li & Schulman[3].

##### 3.4 Annealing protocol

The solutions in Supplemtnary Sections 3.1-3.3. were each annealed by running the thermal ramp program described in Li & Schulman[3]

##### 3.5 Nanopore purification and fluorescent labeling

After thermal annealing (Supplementary Information 3.4), DNA nanopores without adapters were purified using 100kDa Amicon µltra-0.5mL centrifugal filter units (Millipore Sigma UFC510096).The final concentration of the purified nanopores, generally about 1 nM, was measured as described previously[3].

The nanopores that had attached adapters used to assemble nanotubes were purified using the same filter units but were concentrated during the purification process to a final concentration of 2 nM. Specifically, 100 µl of pore solution and 300 µl TAEM buffer were added to a filter unit and centrifuged at 3000 RCF for 4 min in a fixed-angle centrifuge. The sample was washed two more times by adding 300 µl TAEM buffer into the remaining solution and repeating centrifugation. The purified pore solution was then collected by spinning the inverted filter in a new tube.

In both cases, 0.15 µl of 100 µM ATTO647 labeling strand was added to approximately 40 µl purified nanopores collected from the filter unit and was incubated at room temperature for 15 minutes at room temperature.

##### 3.6 Assembly of nanotube channels

To assemble DNA monomers into nanotubes attached to DNA nanopores, DNA monomers were first annealed separately and then mixed with purified nanopores prepared as described in Supplementary Note 3.5.

The anneal was performed by first preparing 20 µl solution containing 400 nM of each of the inactive SEs monomer strands (as listed in Supplementary Note 2) in TAEM buffer and then annealed the mixture using the annealing protocol in Supplementary Note 3.3.

20 µl of the purified nanopores were then mixed with the prepared 20 µl annealed inactive SEs monomers and 0.2 µl of a 50 µM solution of activation strand. The resulting solution was then incubated at 37$^{o}$C for 3-5 hours to allow the nanotubes to grow.

##### 3.7 Hydrophobic modification

To functionalize DNA nanopores with hydrophobic moieties, 1 µl DNA-cholesterol conjugate (“cholesterol strand” in Supplementary Note 1) at 10 µM concentration was added to either 40 µl nanopores without adapters after fluorescent labeling or 40 µl DNA nanotube channels that were assembled on nanopores. The solution was then incubated for 10 minutes at room temperature.

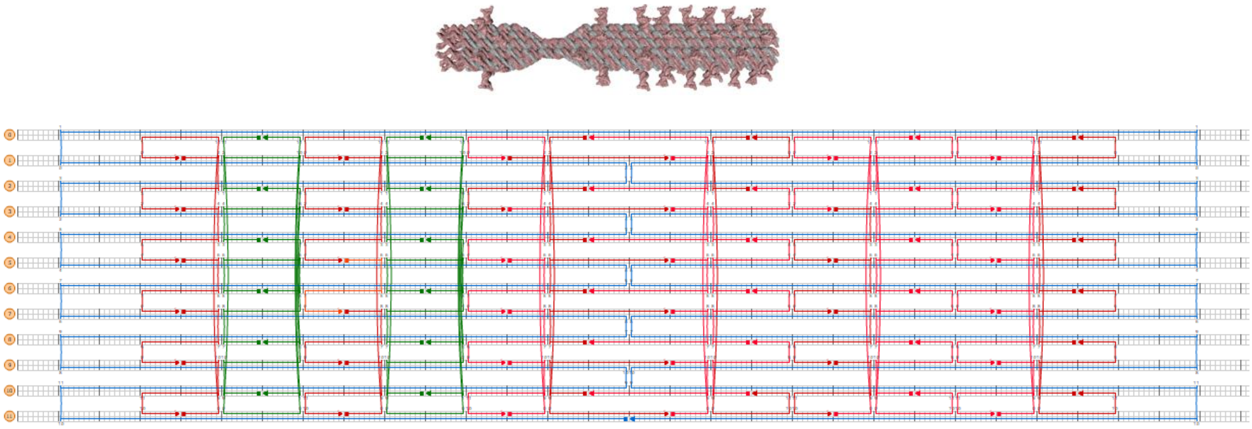

**Supplementary Figure 3. Design of the DNA cap.** Schematic of the DNA cap's structure (The arrangement of its staples on the origami scaffold) produced using CADNano[5] software. The adapters that allow the cap to bind to a nanochannel or a DNA nanopore are not shown here. The staples in red are the as the corresponding staples in the origami pore. The 12 staples in green, whose sequences are listed in Supplementary Table 2, are arranged to create a narrow neck in the structure.

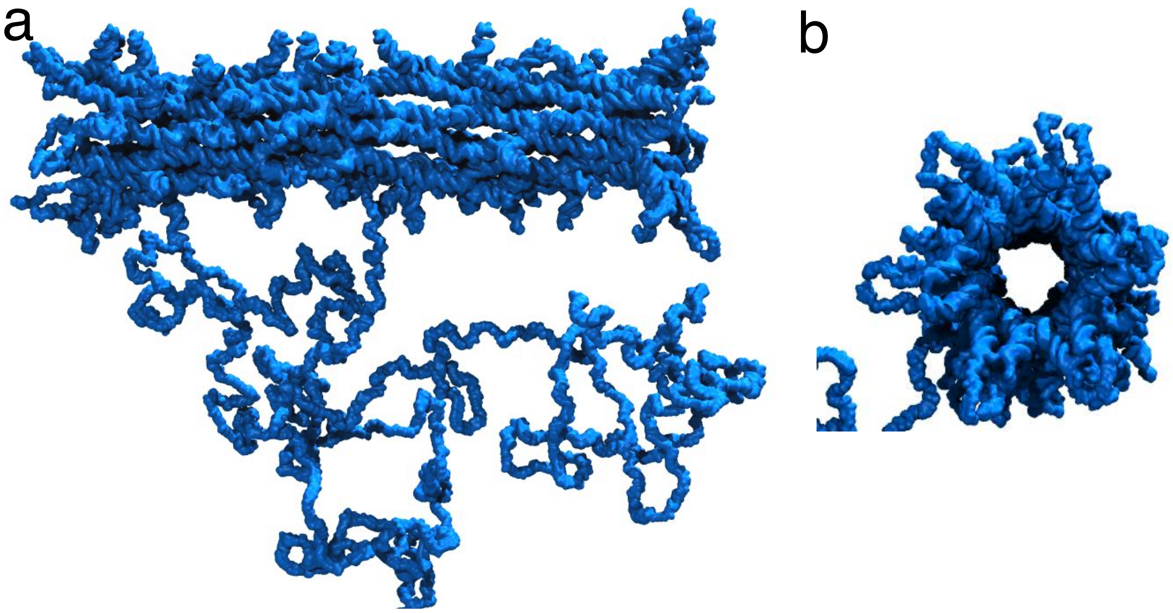


**Supplementary Figure 4. All-atom model of the DNA nanopore structure obtained through multi-resolution simulations. a**) Side view of the cylindrical barrel. **b**) Top view down the axis of the cylinder. Starting from the caDNAno[5] design, the DNA nanopore was simulated using the mrDNA package[6]. In the first 20 μs, the nanostructure was simulated at 1 bead per four base pair resolution, followed by an 8 μs simulation at 1 bead per nucleotide resolution. The final equilibrated was mapped to an all-atom model using mrDNA. The DNA strands are shown using a molecular surface representation. The single-stranded loop below the side view of the pore structure is not part of the nanopore but is used for fluorescence labeling (Supplementary Note 3.5). The predicted inner diameter of the cylinder was determined by averaging the lengths of 40 lines across the cylinder's interior starting at different positions along the helix. The mean inner diameter determined using this method was 7.3 nm ± 0.4 nm.

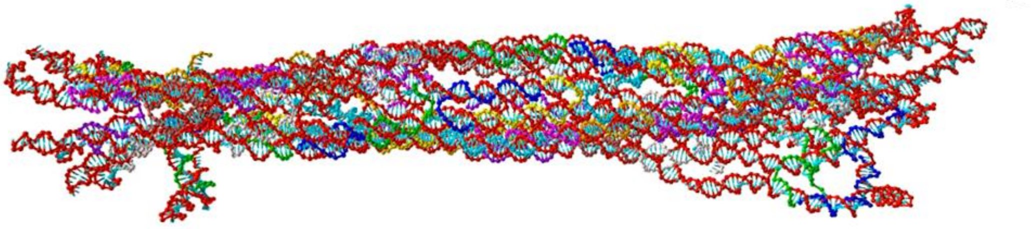

**Supplementary Figure 5. A structural snapshot from a coarse-grained model of the DNA cap.** The model was generated with oxDNA[7] program using an initial configuration consisting of the PDB generated by caDNano (Supplementary Figure 3). The oxDNA scripts for generating the model are available upon request.

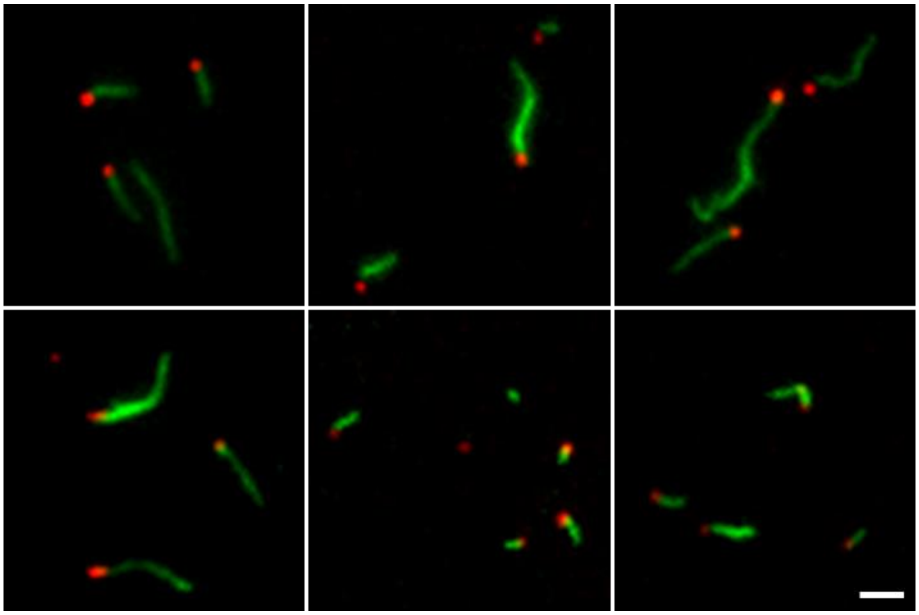

**Supplementary Figure 6. Example fluorescence micrographs of DNA nanotube channels attached to DNA pores**. The nanotubes (Cy3, green) and pores (ATTO647, red) were prepared as described in Supplementary Note 3 except that no DNA-cholesterol conjugate was added. The lengths of the nanotubes were measured by drawing segmented lines along the nanotubes in the images using ImageJ software. Nanotubes were not visible on 13% ± 3% (N = 429) nanopores in the fluorescence images. Scale bar, 2 μm.

**Supplementary Note 4. Preparation of samples for transmission electron microscopy**

Nanostructure samples were deposited on a formvar/carbon film support grid (Cat# FCF400-Cu, Electron Microscopy Sciences, Hatfield, PA, US) to be imaged. To prepare samples of DNA nanopores, nanotube channels, caps, and capped nanopores, 10 µl of the corresponding structures were prepared without attached DNA-cholesterol conjugates in TAEM buffer (Supplementary Note 3), and then were directly used to prepare the grids.

For transmission electron microscopy (TEM) imaging of nanopores on SUVs and nanotube channels on SUVs, SUVs were first prepared and diluted as described in the Methods. The nanopores or nanotube channels were prepared with attached DNA-cholesterol conjugates in TAEM buffer. To prepare nanopores on SUVs, 7.5 µl nanopores were then mixed with 2.5 µl SUVs. To prepare nanotube channels on SUVs, 8 µl nanotube channels were then mixed with 2 µl SUVs. These mixtures were each incubated at room temperature for 10 minutes before use for preparing the grids.

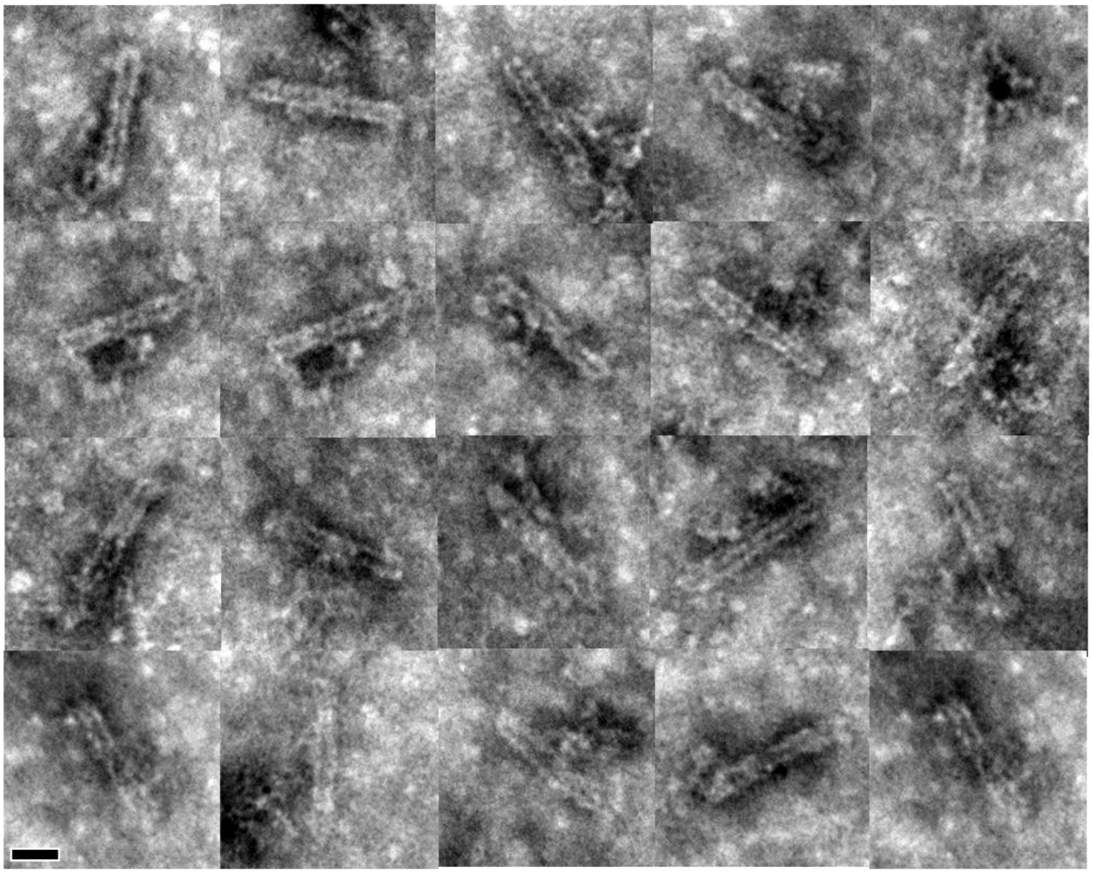

**Supplementary Figure 7. Additional TEM Images of DNA origami nanopores without added DNA-cholesterol conjugates.** Scale bar, 20 nm.

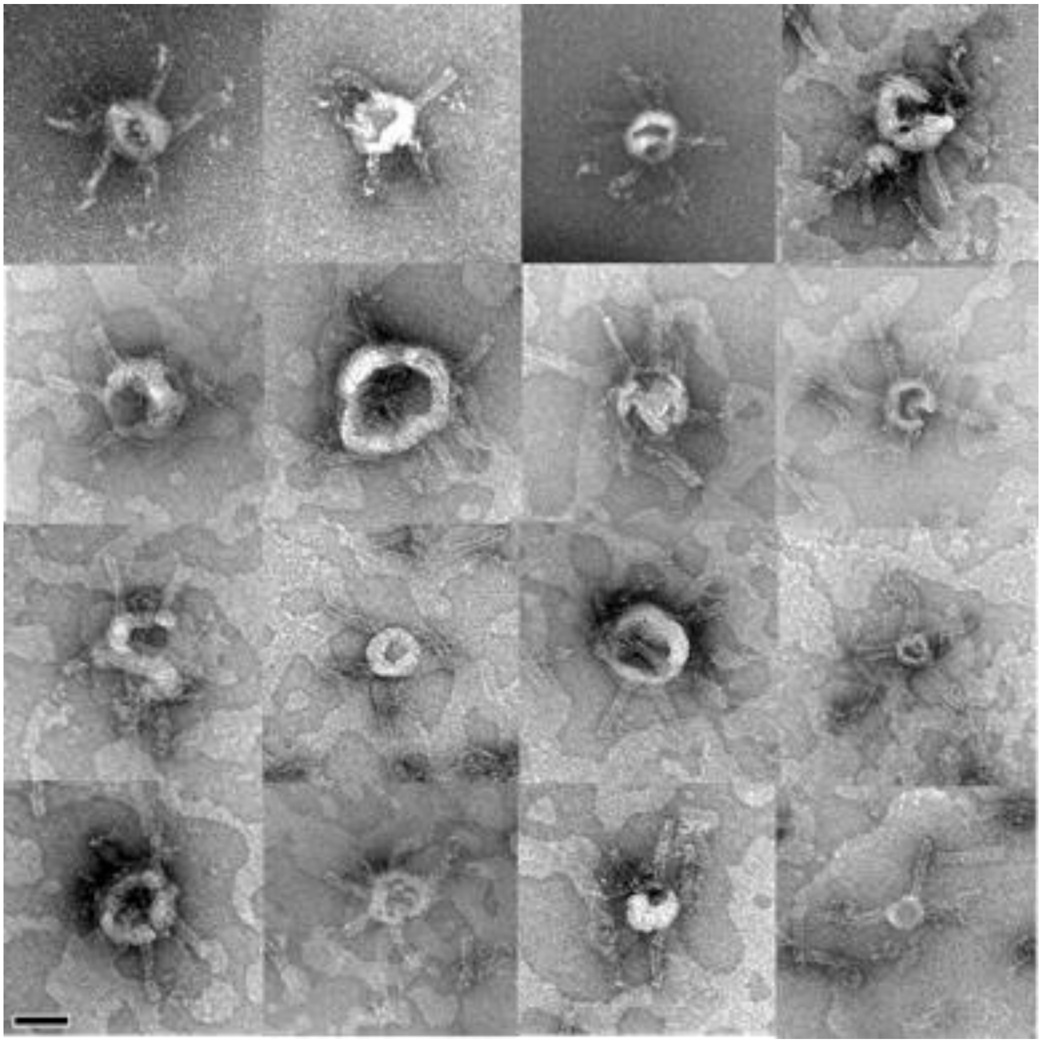

**Supplementary Figure 8. Additional TEM Images of DNA nanopores interacting with SUVs**. Scale bar, 50 nm.

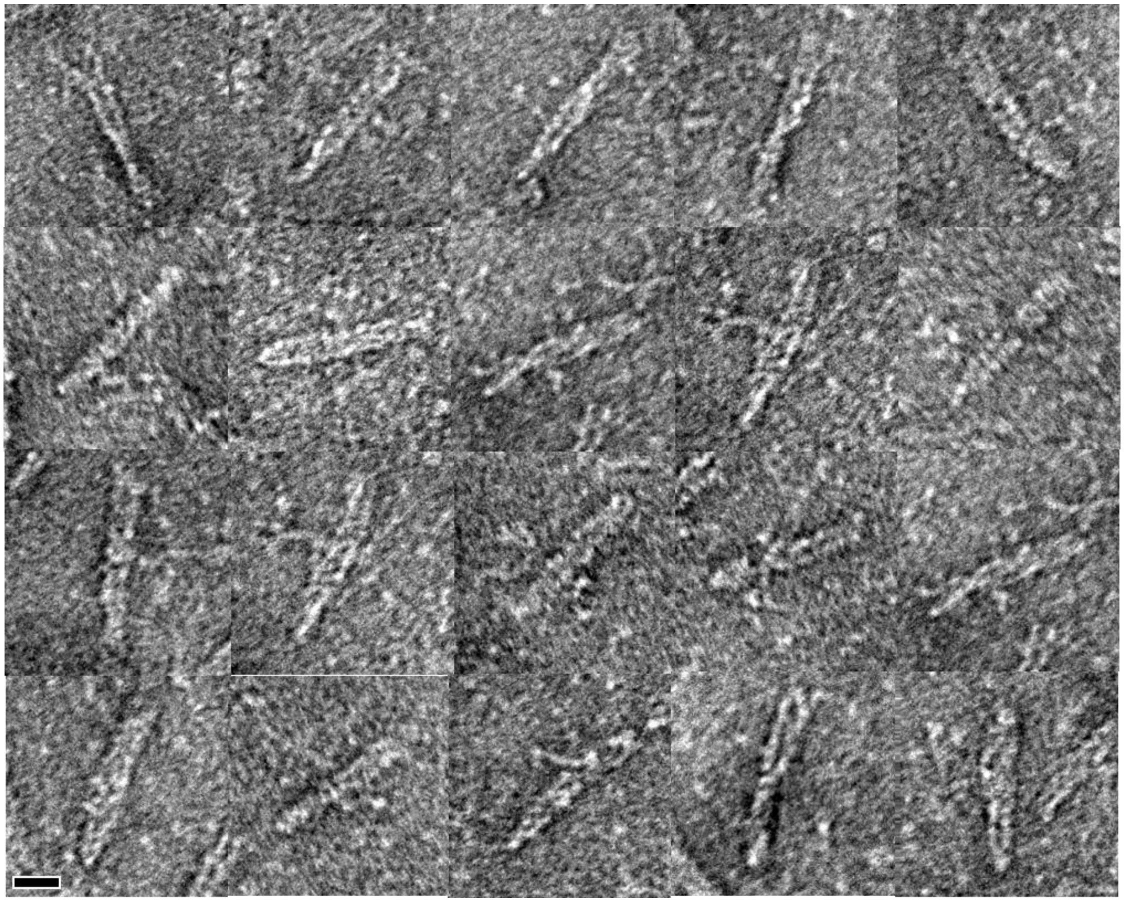

**Supplementary Figure 9. Additional TEM Images of DNA origami caps.** Scale bar, 20 nm.

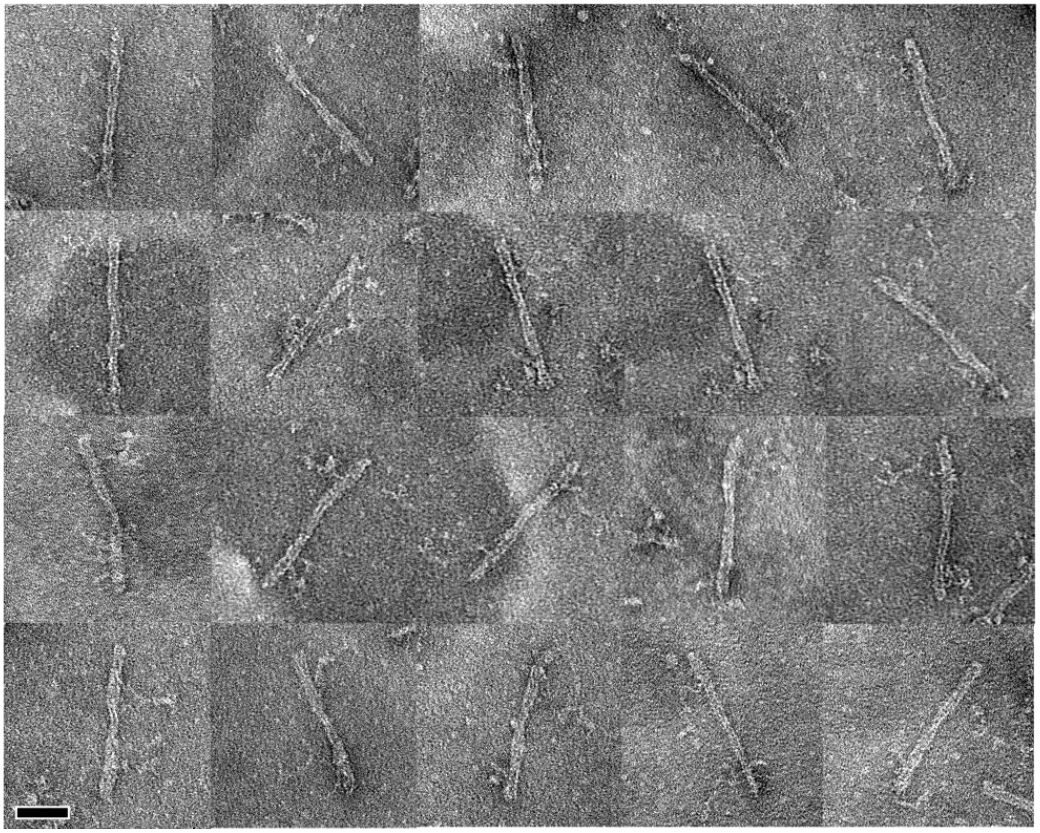

**Supplementary Figure 10. TEM Images of capped nanopores**. The images show how the caps end-to-end bind the nanopores. As DNA caps and pores have similar morphologies, it is difficult to distinguish them in the images. Scale bar, 50 nm.

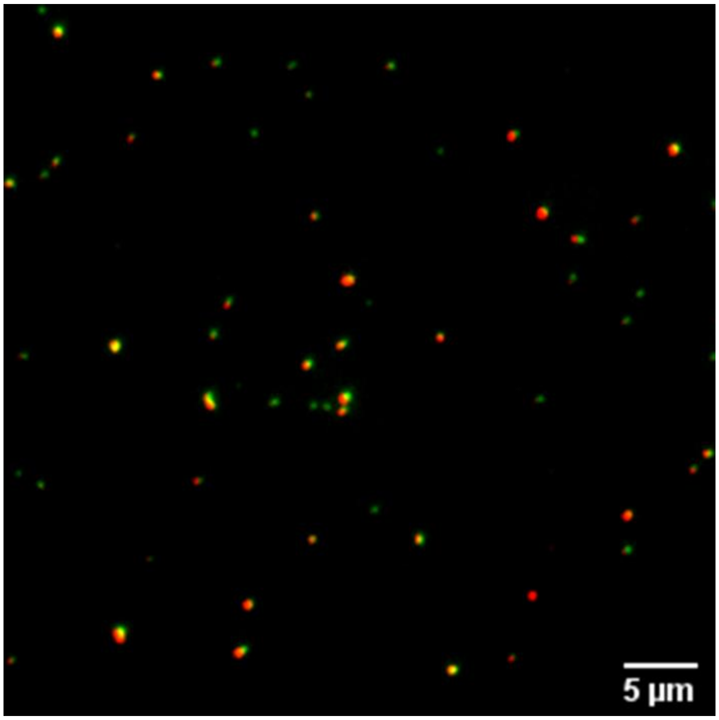


**Supplementary Figure 11. Example wide-field fluorescence image of DNA caps bound to DNA nanopores.** The pores labeled with ATTO647 (red) were mixed with two-fold concentration of caps labeled with ATTO488 (green). The mixture was incubated at room temperature for 3 hours before being imaged on a glass coverslip. The pores were considered capped if the centers of the pores and the caps were within 5 pixels (1 pixel=168 nm). 97.3 ± 0.6% (SD, N=678) pores were capped.

**Supplementary Note 5. A bulk diffusion model of TAMRA influx into vesicles through DNA nanopores**

We model the influx of TAMRA into the vesicles as diffusive transport of TAMRA molecules from a bulk solution into a compartment through DNA nanopores. The concentration gradient of TAMRA between the bulk solution and the compartment drives net diffusion (*i.e.*, influx) of TAMRA into the vesicle. A pore is modeled as a rigid cylindrical channel of diameter *d* and length *L*.

Because the volume of the bulk solution is much larger than the volume of the compartment, the TAMRA concentration in the bulk solution remains constant in the model. Complete influx of TAMRA into vesicles (so that the concentration inside a vesicle and in the bulk are approximately the same) takes half an hour to several hours. The time scale of mixing within the vesicle compartment is therefore much smaller than the time scale of transport from the bulk solution into the compartment. The solution in the compartment can thus be viewed as a uniform bulk phase (quasi-steady-state approximation). This assumption is consistent with our observation in confocal micrographs that fluorescence intensities of TAMRA within the vesicles do not show spatial variations.

The molar flux of TAMRA (*J*) through the channels into a vesicle is given by

$$J = D\frac{C_{out}-C_{in}}{L} \qquad (1)$$

where *D* is the diffusion coefficient of TAMRA and $C_{out}$ and $C_{in}$ are bulk phase concentrations of TAMRA in the bulk phase outside and inside the compartment respectively.

The molar flow rate of TAMRA into a vesicle can be written as

$$\dot{n} = J \cdot A = \frac{AD}{L}(C_{out} - C_{in}) \qquad (2)$$

where *A* is the total of the cross-sectional areas of the channels spanning the membrane. The change in the amount of TAMRA inside the compartment is then given by

$$\frac{dN}{dt} = \dot{n} = V\frac{dC_{in}}{dt} \qquad (3)$$

where N is amount (in moles) of TAMRA inside the vesicle and *V* is the volume of the vesicle. Equating (2) and (3) gives

$$\frac{dC_{in}}{dt} = \frac{AD}{VL}(C_{out} - C_{in}) \qquad (4)$$

Rearranging the equation to separate variables then gives

$$\frac{dC_{in}}{C_{out}-C_{in}} = \frac{AD}{VL}dt \qquad (5)$$

We now introduce the fractional concentration $f = \frac{C_{in}}{C_{out}}$, defined as the ratio of the TAMRA concentration inside the vesicle to the concentration outside. Substituting $f$ into equation (6) gives

$$\frac{df}{1-f} = \frac{AD}{VL}dt \qquad (6)$$

We can then solve for fractional concentration as a function of time by integrating the differential equation with the initial condition of $f(t=t_1) = f_1$ where $t_1$ is the time when the influx starts:

$$f_{pore} = f_1 + (1 - f_1)[1 - e^{-\frac{(t-t_1)}{\tau}}] \qquad (7)$$

Here we introduce the abbreviation, time constant $\tau = \frac{VL}{AD}$, which has unit of minutes. Equation (7) describes how fractional TAMRA intensity changes due to TAMRA influx through the pores, so this fractional influx is denoted as $f_{pore}$.

"Leak" transport of TAMRA across the membrane (*i.e.* transport not mediated by nanopores) also occurs at rates described by equation (7). However, the leaky transport rate is much slower than the pore-mediated transport as observed in the dye influx experiments, so the time constant $\tau$ for the leaky transport is very small. We use a linear approximation for the kinetics t during both the lag time and the influx phase, written as

$$f_{leak} = f_0 + a * t \qquad (8)$$

where $a$ is the leaky influx kinetic parameter of TAMRA into a vesicle.

Before a nanopore or nanopores insert into a vesicle ($t < t_1$), the fractional concentration increases solely because of leak transport. During this period, the fractional concentration as a function of time is therefore

$$f(t) = f_0 + a * t \qquad \text{when } t < t_1. \qquad (9)$$

The fractional intensity when influx starts is therefore

$$f_1 = f_0 + a * t_1 \qquad (10)$$

After nanopore insertion ($t=t_1$), increases in the fractional concentration are attributed to both leak transport and pore-mediated influx. The fractional concentration as a function of time is therefore

$$f = f_1 + a(t - t_1) + (1 - f_1)\left[1 - e^{-\frac{(t-t_1)}{\tau}}\right] \qquad \text{when } t \geq t_1. \qquad (11)$$

Combining the two fractional concentration functions and plugging in the equation for $f_1$ results in a piecewise function for the fractional concentration:

$$f = \begin{cases} f_0 + a * t & \text{when } t < t_1 \\ f_0 + a * t + (1 - f_0 - ct_1)\left[1 - e^{-\frac{(t-t_1)}{\tau}}\right] & \text{when } t \geq t_1 \end{cases} \quad (12)$$

#### Supplementary Note 6. Rate of TAMRA influx through a single DNA nanopore

The influx rate ($k$) of TAMRA into a vesicle through a single nanopore measured to be 13.1 ± 1.5 μm$^3$/min in experiments (see main text). To convert this rate into units of molecules per second, we plug $k = \frac{V}{\tau} = \frac{AD}{L}$ into Equation 3 and 4, and rearrange to write

$$\frac{dN}{dt} = V\frac{dC_{in}}{dt} = k[C_{out} - C_{in}(t)] \qquad (13)$$

Equation (13) can be used to calculate the influx rate in the unit of molecules per second for each vesicle at a time (*t*) after influx starts. By inserting $k$ = 13.1 ± 1.5 μm$^3$/min, $C_{out}$ = 309 nM, and $C_{in}$=0 into Equation (13), we find that at t = 0, the flux of TAMRA through a single nanopore should be 40.5 ± 4.6 molecules per second.

**Supplementary Note 7. A bulk diffusion model of TAMRA influx into vesicles through DNA nanochannels**

The DNA nanochannels are cylindrical structures that have the same diameters as the DNA pores but longer lengths. We model influx of TAMRA through nanotube channels using the same model used for transport through DNA pores but adjust the channel length parameter. In this case, the fractional concentration of a vesicle as a function of time can be written as in Eq 12.

For nanotube-mediated transport, the channel-mediated transport rate $k = \frac{V}{\tau} = \frac{AD}{L}$, should in general be smaller than the channel-mediated transport rates in for nanopore-mediated transport due to longer channel lengths ($L$).

Eq 12 is written to assume a single influx event but could be extended to account for multiple insertion events. Accounting for multiple insertion events would increase the complexity of fitting but would be required to properly fit influx curves in which there were multiple insertion events at different times. We had sufficient data from experiments with DNA nanopores to measure the distribution of influx rates and to deduce the influx rate through a single pore using only traces with single influx events (Main text and Supplementary Note 10). But experiments with DNA nanochannels produced fewer traces with single influx to characterize rates of transport through open and capped nanotube channels. To measure the influx rates for eight vesicles in the dye influx experiments with the DNA nanotube channels that showed two distinct curves, we modified the diffusion model described in Supplementary Note 5 to account for influx events with two sequential insertions.

In the case of two sequential insertions, the fractional concentration as a function of time, *f(t)*, follows Eq 12 until an additional influx event starts at $t=t_2$, *i.e.*:

$$f = \begin{cases} f_0 + a * t & \text{when } \mathrm{t} < \mathrm{t}_1 \\ f_0 + a * t + (1 - f_0 - a * t_1)\left[1 - e^{-\frac{(t-t_1)}{\tau_1}}\right] & \text{when } \mathrm{t}_1 \le \mathrm{t} < \mathrm{t}_2 \end{cases} \qquad (14)$$

where $\tau_1$ is the influx time constant for the nanotube channels inserted at $t = t_1$. The influx rate after $t = t_2$ is then the sum of the influx rates due to leak transport, pore-mediated influx that starts at $t = t_1$, and pore-mediated influx that starts at $t = t_2$. Thus, the fractional concentration in the case where there are two insertions of a nanotube channel channels, at $t_1$ and $t_2$ respectively is

$$f = \begin{cases} f_0 + a * t & \text{when } \mathrm{t} < \mathrm{t}_1 \\ f_0 + a * t + (1 - f_0 - a * t_1)\left[1 - e^{-\frac{(t-t_1)}{\tau_1}}\right] & \text{when } \mathrm{t}_1 \le \mathrm{t} \le \mathrm{t}_2 \\ f_0 + a * t + (1 - f_0 - a * t_1)\left[1 - e^{-\frac{(t-t_1)}{\tau_1}}\right] + (1 - f_2)\left[1 - e^{-\frac{(t-t_2)}{\tau_2}}\right] & \text{when } \mathrm{t} > \mathrm{t}_2 \end{cases} \qquad (15)$$

where $\tau_2$ is the influx time constant for the nanotube channels inserted at $t = t_2$ and $f_2$ is the fractional concentration at $t=t_2$:

**Supplementary Note 8. Image analysis of fluorescence micrographs from dye influx experiments**

In order to measure the changes in fluorescence intensities of hundreds of GUVs using fluorescence images captured over time during influx experiments, an image analysis algorithm was developed using ImageJ (version 2.1.0/1.53c) and MATLAB (version 2020a) software.

The first step of the algorithm was to locate the vesicles and to determine their respective volumes. The 8-bit grayscale images of the vesicle membrane fluorescence channel and the TAMRA fluorescence channel were imported into ImageJ software as two time series stacks. The image stack of the vesicle membrane channel was converted into a binary image stack by taking the threshold of 20 pixel-intensity units to find the outlines of vesicles in the images (Supplementary Figure 10). The "Analyze particles" function in ImageJ was then applied to the binary image stack to find all the circular objects (vesicles) with diameters over 4 µm (corresponding to actual vesicle diameter of 5 µm, as explained below) and extents of circularity at least 0.6 in the stack.

Because the 2-D circular cross-sections of the vesicles that appeared in the confocal fluorescence images captured at the specific height were not necessarily the largest cross-sections of the vesicles, the diameters of the vesicle cross-sections in the images needed to be converted to the vesicles' actual diameters (Supplementary Note 9). Because a 4 µm vesicle diameter in the confocal images corresponded to an actual vesicle diameter of 5 µm (Eq 17), a 4 µm diameter limit was used in the search criteria in the image processing algorithm.

The "Analyze particles" function then measured and recorded the sizes of coordinates of the vesicles that fit the searching criteria. Meanwhile, the mean fluorescence intensities of TAMRA both inside and outside the vesicles were measured in the TAMRA fluorescence channel using the vesicle outlines found. The fluorescence intensities of TAMRA inside each vesicle and outside of the vesicles were then used to determine the fractional concentrations of TAMRA inside each vesicle over time.

Because the vesicles were immobilized to the surface by biotin-streptavidin linkages, the vesicles showed minimal movement over time. Thus, each vesicle found in each image in the stack was matched with the same vesicle across the stack based on its coordinates and size, so that the interior mean TAMRA intensities over time for each vesicle were obtained. The fractional intensity of each vesicle at each time point was calculated by dividing the interior intensity by the exterior intensity at each time point. We excluded vesicles that burst or ruptured during the experiment by removing the vesicles that had an initial fractional intensity less than 0.5, an increase of over 0.1 in the fractional intensity within time interval of 1 minute, or were observed in fewer than 60% of total time points.

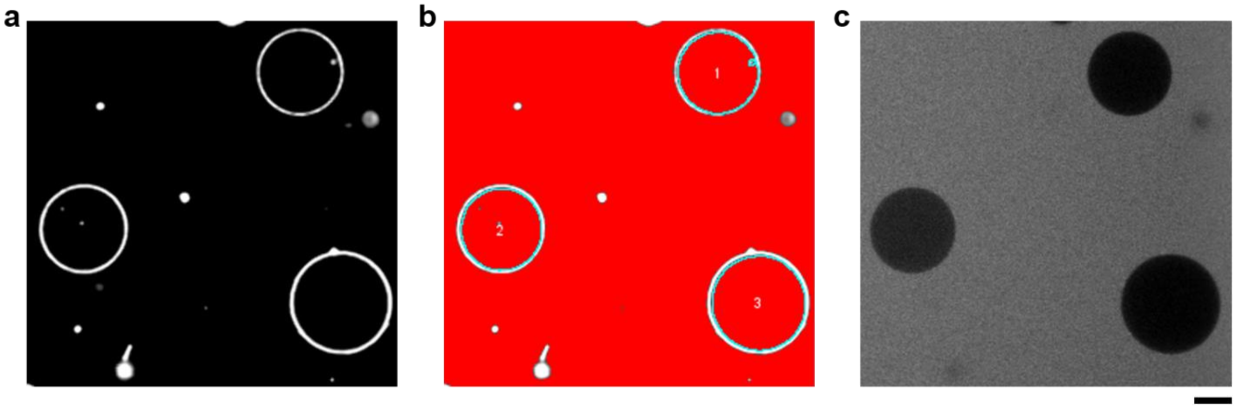


**Supplementary Figure 12. Example fluorescence micrographs of GUVs from the dye influx experiment with DNA nanopores.** a) Confocal fluorescence images in the vesicle membrane fluorescence channel were used to determine the vesicle locations within the images and their volumes. b) A binary image (red areas are retained after thresholding), who's on points contained the areas of the vesicles, was generated by using a threshold filter on the image in (a) that selected pixels with intensity values in the range of 0 – 20 pixel intensity values. The perimeter of each circle in the image was used to calculate the volume of the vesicle by assuming each vesicle had a spherical shape and using Equation 17 (Supplementary Note 9) to convert measured diameter to actual vesicle diameter. The fluorescence intensities of TAMRA inside and outside the vesicles were measured in c, the grayscale fluorescence micrograph in the TAMRA channel, using the image areas determined in b). Scale bar, 10 µm.

**Supplementary Note 9. Calculating vesicle volumes in the dye influx experiments**

To quantify the influx rates in the dye influx experiments from the changes in fractional concentration inside a vesicle, we needed to determine the volume of each vesicle. The confocal fluorescence images in the vesicle membrane channel show two-dimensional cross-sections of the vesicles (Supplementary Figure 10), which have a circular shape. The perimeters of these circles were measured using ImageJ software.

The measured radius of a circle, *w*, was converted to the vesicle's spherical radius, *R*, by

$$R = \frac{(w^2+z^2)}{2z} \qquad (16)$$

where *z* is the height of the focal distance from the coverslip surface, which was set to 8 μm in all dye influx experiments. The measured vesicle volumes changed during the experiments, which were mostly fluctuating throughout time-lapsed imaging while some vesicles showed increasing sizes over time. To account for these measurement variations, for each vesicle, we calculated the mean volume and associated standard errors across all time points. The mean volume and standard errors were then used in obtaining the Gaussian distribution of the influx rate (Supplementary Note 10).

**Supplementary Note 10. Regression analysis of fractional intensity data**

To determine influx rates into each vesicle using Equation 12, we first converted the time-lapse measurements of fractional intensities into piecewise fractional concentrations using the assumption that the measured fluorescence intensity is proportional to the dye concentration.

We performed regression analysis using the diffusion models developed in Supplementary Notes 5 and 7 to quantify the four kinetic parameters that described influx kinetics for each vesicle. These parameters are 1) the initial fractional intensity ($f_0$), 2) the linear leaky transport rate ($k_0$), 3) the time at which fast influx starts ($t_1$), and 4) the influx time constant ($\tau$). For vesicles that experienced two fast influx events in the nanotube channel dye influx experiment, two additional parameters, the second influx time constant ($\tau_2$) and the time when the second influx starts ($t_2$), were also fit.

The regression was performed by using the nonlinear least-squares solver (“lsqcurvefit”) in MATLAB to find the four (or six) parameters. The 95% confidence interval of $\tau$ during the regression were calculated using the “nlparci” function in MATLAB, from which the standard error of $\tau$ for each vesicle was calculated. The standard errors in vesicle volumes ($V$) were also calculated from the measurements of vesicle diameters in the confocal image. The values of the fast influx rate ($k_f = \frac{V}{\tau}$) and the second fast influx rate ($k_{f2} = \frac{V}{\tau_2}$) for the corresponding vesicles were then calculated. Finally, the standard errors and 95% confidence intervals of $k$ and $k_2$ were calculated through propagation of error.

To obtain the distribution of influx rates for the vesicles that accounts for the uncertainties in fit, a Gaussian distribution function was fitted to each influx rate using the calculated influx rate and standard error. The probability density function of influx rates of all measured vesicles was obtained by summing the Gaussian distribution of each influx rate in the range of 0 to 120 $\mu m^3$/min and normalizing the integral to 1. The cumulative distribution function was calculated by numerically integrating the probability density function of the influx rates.

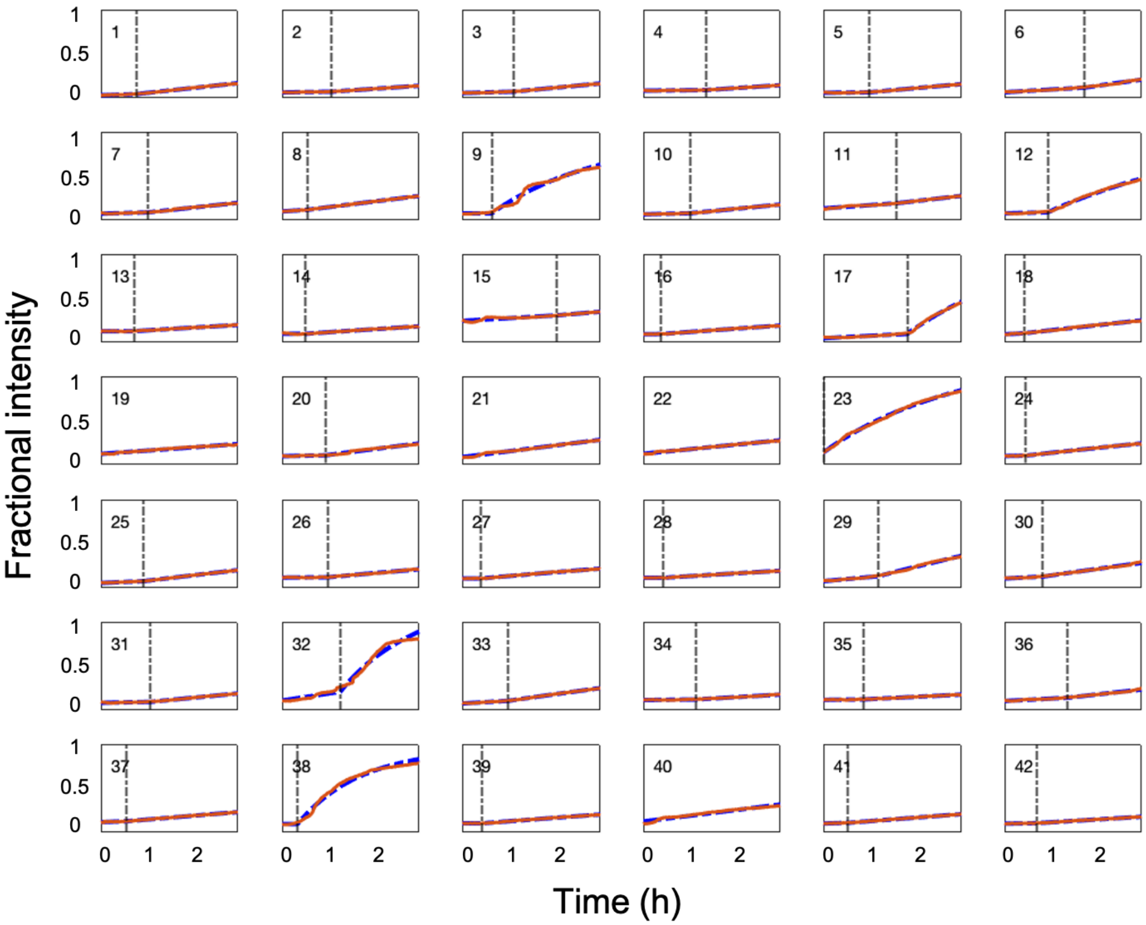


**Supplementary Figure 13. Example plots of measured fractional intensities of GUVs and corresponding regression curves**. The red curve is the measured fractional intensities of tracked GUVs in the dye influx experiment with DNA nanopores. The blue dash curve is the regression curve by fitting the diffusion model to the measurements (Supplementary Note 10). The vertical dash-dotted lines in the plots, in the plots for vesicles that experienced fast influx, indicate the times when fast influx starts ($t_1$), obtained through regression. The plots are ordered by decreasing vesicle volume, and vesicle numbers are shown inside the plots (number 1 is the plot of the largest vesicle). The influx rates, $k$, and the times at which fast influx of TAMRA starts, $t_1$, are determined through regression. The volumes of the GUVs, $V$, are measured in the confocal images during the dye influx experiment (Supplementary Note 9). These parameters for the vesicles in the figure are listed in Supplementary Table 1.

**Supplementary Table 3. Kinetic parameters for vesicles in Supplementary Figure 13.**

| Vesicle Number | $a\ (min^{-1})$ | $f_0$ | $t_1\ (min)$ | $V\ (\mu m^3)$ | $k_f\ (\frac{\mu m^3}{min})$ |
|---|---|---|---|---|---|
| 1 | 2.63E-04 | 0.032 | 51.10 | 2.65E+04 | 18.13 |
| 2 | 1.26E-04 | 0.066 | 71.84 | 2.53E+04 | 12.10 |
| 3 | 2.30E-04 | 0.058 | 73.42 | 2.34E+04 | 14.29 |
| 4 | 8.42E-05 | 0.087 | 90.00 | 1.79E+04 | 9.15 |
| 5 | 7.95E-05 | 0.062 | 65.72 | 1.69E+04 | 12.48 |
| 6 | 4.82E-04 | 0.068 | 115.93 | 1.66E+04 | 13.09 |
| 7 | 2.22E-04 | 0.078 | 68.39 | 1.59E+04 | 12.07 |
| 8 | 5.43E-04 | 0.104 | 36.86 | 1.55E+04 | 8.90 |
| 9 | 2.22E-14 | 0.082 | 42.20 | 1.28E+04 | 81.82 |
| 10 | 1.90E-04 | 0.069 | 67.66 | 1.25E+04 | 8.62 |
| 11 | 5.85E-04 | 0.140 | 104.03 | 1.20E+04 | 5.53 |
| 12 | 1.99E-04 | 0.081 | 62.37 | 1.19E+04 | 46.89 |
| 13 | 2.35E-14 | 0.133 | 47.65 | 1.17E+04 | 6.99 |
| 14 | 2.22E-14 | 0.103 | 33.86 | 1.17E+04 | 7.41 |
| 15 | 4.82E-04 | 0.256 | 135.49 | 1.13E+04 | 3.65 |
| 16 | 3.27E-05 | 0.098 | 25.02 | 1.12E+04 | 7.33 |
| 17 | 4.23E-04 | 0.050 | 119.91 | 1.11E+04 | 69.91 |
| 18 | 4.38E-04 | 0.096 | 27.88 | 1.06E+04 | 5.85 |
| 19 | 5.96E-04 | 0.127 | 1.00 | 1.01E+04 | 0.00 |
| 20 | 1.73E-04 | 0.100 | 63.35 | 9.99E+03 | 10.43 |
| 21 | 1.04E-03 | 0.089 | 1.03 | 9.90E+03 | 0.00 |
| 22 | 8.19E-04 | 0.129 | 4.58 | 9.82E+03 | 0.00 |
| 23 | 6.38E-04 | 0.148 | 1.00 | 9.36E+03 | 60.21 |
| 24 | 1.16E-04 | 0.103 | 29.99 | 9.09E+03 | 8.32 |
| 25 | 3.42E-04 | 0.059 | 61.90 | 8.70E+03 | 6.20 |
| 26 | 5.85E-05 | 0.123 | 65.82 | 8.55E+03 | 6.85 |
| 27 | 2.22E-14 | 0.113 | 26.48 | 8.54E+03 | 7.10 |
| 28 | 1.04E-05 | 0.121 | 29.19 | 8.06E+03 | 4.70 |
| 29 | 7.40E-04 | 0.081 | 78.04 | 7.69E+03 | 12.69 |
| 30 | 4.04E-04 | 0.113 | 54.81 | 7.67E+03 | 7.04 |
| 31 | 1.47E-04 | 0.088 | 71.24 | 7.55E+03 | 5.62 |
| 32 | 1.25E-03 | 0.114 | 83.79 | 7.55E+03 | 87.62 |
| 33 | 4.77E-04 | 0.079 | 65.14 | 7.43E+03 | 5.90 |
| 34 | 8.64E-05 | 0.121 | 74.79 | 7.36E+03 | 3.38 |
| 35 | 8.14E-05 | 0.123 | 57.13 | 7.13E+03 | 2.77 |
| 36 | 4.20E-04 | 0.109 | 90.07 | 6.95E+03 | 4.10 |
| 37 | 2.83E-04 | 0.122 | 36.87 | 6.82E+03 | 3.24 |
| 38 | 2.22E-14 | 0.099 | 22.35 | 6.79E+03 | 75.19 |
| 39 | 2.22E-14 | 0.107 | 28.34 | 6.59E+03 | 4.90 |
| 40 | 1.03E-03 | 0.130 | 1.00 | 6.56E+03 | 0.00 |
| 41 | 2.23E-04 | 0.105 | 33.88 | 6.52E+03 | 3.17 |
| 42 | 1.26E-04 | 0.103 | 45.85 | 6.27E+03 | 2.76 |

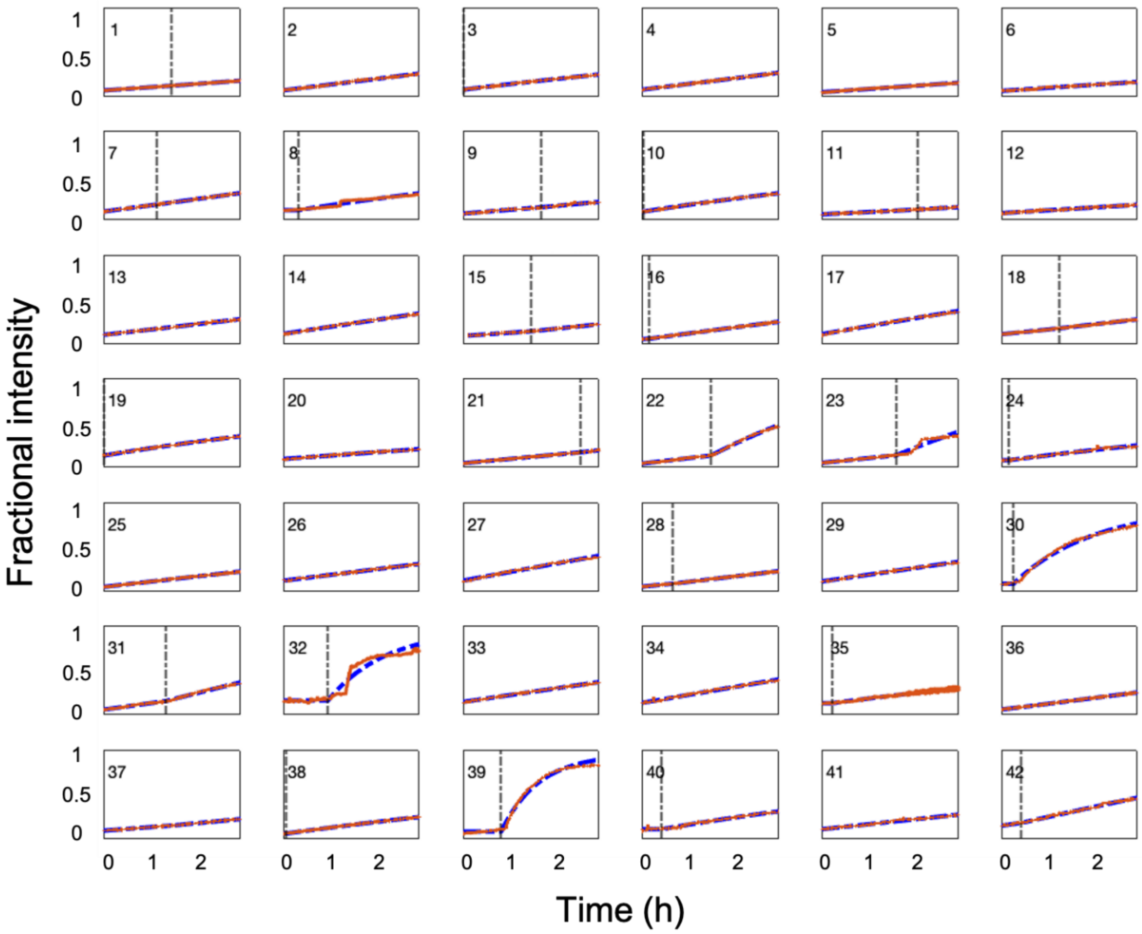


**Supplementary Figure 14. Example plots of measured and fitted fractional intensities of GUVs in experiments where DNA nanochannels are added to GUVs.** The red curve in each plot is the measured fractional intensity of a tracked GUVs. The blue dash curve is the regression curve produced by fitting the diffusion model to the measurements (Eq 14 or 15, Supplementary Note 10). The vertical dash-dotted lines, in the plots for vesicles that experienced fast influx, indicate the time when the fast influx starts ($t_0$), obtained through regression. The influx rates, $k$, and the times at which fast influx of TAMRA starts, $t_0$, are determined through regression. The volumes of the GUVs, $V$, are measured in the confocal images during the dye influx experiment (Supplementary Note 9). These parameters for the vesicles in the figure are listed in Supplementary Table 2.

**Supplementary Table 4. Kinetic parameters for vesicles in Supplementary Figure 14.**

| Vesicle Number | $a\ (min^{-1})$ | $f_0$ | $t_1\ (min)$ | $V\ (\mu m^3)$ | $k_f\ (\frac{\mu m^3}{min})$ |
|---|---|---|---|---|---|
| 1 | 3.57E-04 | 0.074 | 156 | 4.09E+05 | 8.53 |
| 2 | 4.41E-04 | 0.076 | 94 | 1.33E+05 | 0.02 |
| 3 | 5.74E-06 | 0.084 | 1 | 1.14E+05 | 54.34 |
| 4 | 4.45E-04 | 0.084 | 136 | 7.78E+04 | 0.01 |
| 5 | 3.03E-04 | 0.054 | 367 | 7.14E+04 | 0.00 |
| 6 | 2.38E-04 | 0.069 | 447 | 5.81E+04 | 0.00 |
| 7 | 4.71E-04 | 0.099 | 173 | 5.02E+04 | 1.52 |
| 8 | 2.22E-14 | 0.115 | 40 | 4.93E+04 | 38.50 |
| 9 | 2.87E-04 | 0.075 | 258 | 4.92E+04 | 2.48 |
| 10 | 2.89E-14 | 0.098 | 5 | 3.73E+04 | 23.15 |
| 11 | 1.79E-04 | 0.066 | 314 | 3.59E+04 | 1.43 |
| 12 | 2.18E-04 | 0.080 | 439 | 3.37E+04 | 24.90 |
| 13 | 4.14E-04 | 0.110 | 219 | 3.17E+04 | 0.39 |
| 14 | 5.34E-04 | 0.123 | 62 | 3.15E+04 | 0.01 |
| 15 | 2.44E-04 | 0.097 | 224 | 2.19E+04 | 3.29 |
| 16 | 2.26E-04 | 0.059 | 24 | 1.87E+04 | 5.26 |
| 17 | 6.14E-04 | 0.118 | 2 | 1.83E+04 | 0.00 |
| 18 | 4.74E-04 | 0.117 | 147 | 1.61E+04 | 1.12 |
| 19 | 1.09E-06 | 0.140 | 3 | 1.60E+04 | 10.92 |
| 20 | 2.60E-04 | 0.099 | 447 | 1.59E+04 | 0.00 |
| 21 | 3.32E-04 | 0.049 | 386 | 1.57E+04 | 1.31 |
| 22 | 4.17E-04 | 0.045 | 225 | 1.55E+04 | 26.14 |
| 23 | 3.65E-04 | 0.053 | 243 | 1.50E+04 | 19.94 |
| 24 | 2.22E-14 | 0.084 | 23 | 1.31E+04 | 6.56 |
| 25 | 4.07E-04 | 0.056 | 1 | 1.21E+04 | 0.00 |
| 26 | 4.37E-04 | 0.129 | 167 | 1.18E+04 | 0.07 |
| 27 | 6.54E-04 | 0.130 | 2 | 1.10E+04 | 0.00 |
| 28 | 3.35E-04 | 0.059 | 99 | 1.03E+04 | 1.08 |
| 29 | 5.17E-04 | 0.120 | 21 | 9.07E+03 | 0.00 |
| 30 | 2.22E-14 | 0.092 | 40 | 8.40E+03 | 31.75 |
| 31 | 4.98E-04 | 0.060 | 202 | 7.71E+03 | 3.97 |
| 32 | 2.22E-14 | 0.168 | 129 | 7.42E+03 | 45.08 |
| 33 | 5.23E-04 | 0.151 | 1 | 7.37E+03 | 0.00 |
| 34 | 6.19E-04 | 0.140 | 4 | 7.36E+03 | 0.00 |
| 35 | 2.22E-14 | 0.137 | 18 | 6.49E+03 | 6.89 |
| 36 | 4.23E-04 | 0.062 | 91 | 5.93E+03 | 0.22 |
| 37 | 2.72E-04 | 0.099 | 231 | 5.64E+03 | 0.47 |
| 38 | 2.22E-14 | 0.066 | 11 | 5.27E+03 | 2.80 |
| 39 | 2.22E-14 | 0.087 | 125 | 5.06E+03 | 41.35 |
| 40 | 9.35E-14 | 0.116 | 64 | 4.72E+03 | 3.34 |
| 41 | 3.86E-04 | 0.115 | 91 | 4.70E+03 | 0.00 |
| 42 | 5.06E-04 | 0.157 | 66 | 4.69E+03 | 1.68 |

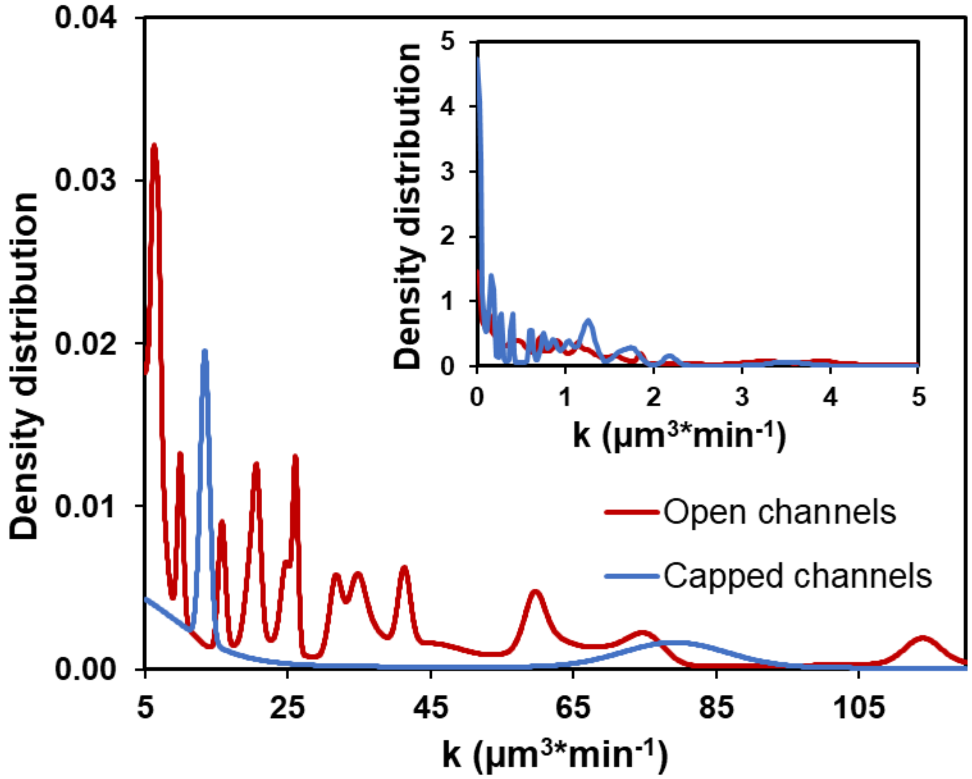


**Supplementary Figure 15. Measured density distributions of the influx rates in the nanochannel and capped nanochannel dye influx experiments**. The influx rates were determined by fitting a diffusion model (either Eq 14 or 15) to the measured influx kinetics of GUVs tracked in the experiments in which DNA channels (N = 138) and capped channels (N = 82) used, respectively, and the ones that fit poorly (SSE > 0.1) were excluded. The influx rates for each GUV were then fit to Gaussian distributions (Supplementary Note 10).

**Supplementary Note 11. Theoretical rate of TAMRA transport through a single DNA pore**

We hypothesize that the rate of one-dimensional transport of TAMRA within the DNA pore that spans across the vesicle membrane follows Fick's laws of diffusion without significant effects due to transport along the pore surface. In this case, the net flux of TAMRA transport through a single DNA pore is described by Eq 2. Because TAMRA concentration inside the vesicle ($C_{in}$) increases over time during the influx event, we define the pore mediated fast influx rate

$$k_f = \frac{V}{\tau} = \frac{AD}{L} \tag{18}$$

as a measurement of how fast TAMRA transports through the pores. Here, $\tau$ is the time constant of the negative exponential kinetics, $V$ is the volume of the vesicle, $A$ is the cross-sectional area of the pore, $L$ is the length of the pore, and $D$ is the diffusion coefficient of TAMRA.

To calculate the theoretical influx rate for a single pore, we calculated the cross-sectional area to be $A = 42 \pm 3.4$ nm$^2$ after approximating the cross-section as a circle with a measured diameter of 7.3 ± 0.4 nm (Supplementary Fig. 2), and the length of the pore was measured $L = 61.1 \pm 2.1$ nm.

In the dye influx experiments, TAMRA was in a 0.2 M glucose solution at 37$^o$C. The diffusion coefficient of TAMRA in such conditions can be calculated from the diffusion coefficient of TAMRA, $D_1 = 2.8 \pm 0.3 * 10^{-6}$ cm$^2$/s, measured in 10% glycerol at room temperature9 based on the Stoke-Einstein Equation, which predicts

$$\frac{D_1}{D_2} = \frac{T_1 \mu_2}{T_2 \mu_1} \tag{19}$$

which accounts for the diffusion coefficient's dependences on temperature and viscosity. Here, D1 and D2 are the diffusion coefficients in the two conditions. T1 and T2 are the corresponding absolute temperatures. $\mu_1$ and $\mu_2$ are the corresponding dynamic viscosities. The dynamic viscosity of 10% glycerol at 37$^o$C is $\mu_1$=9.30*$10^{-4}$ Pa*s.[9] The dynamic viscosity of 0.2 M glucose solution at 37$^o$C is $\mu_2$=7.28*$10^{-4}$ Pa*s after interpolation.[10] The diffusion coefficient of TAMRA in the experimental conditions is then calculated to be $D_2$=3.58 ± 0.4 *$10^{-6}$ cm$^2$/s. Using this value, we find the theoretical influx rate of a single DNA pore is $k_f$ = 14.7 ± 1.8 μm$^3$/min.

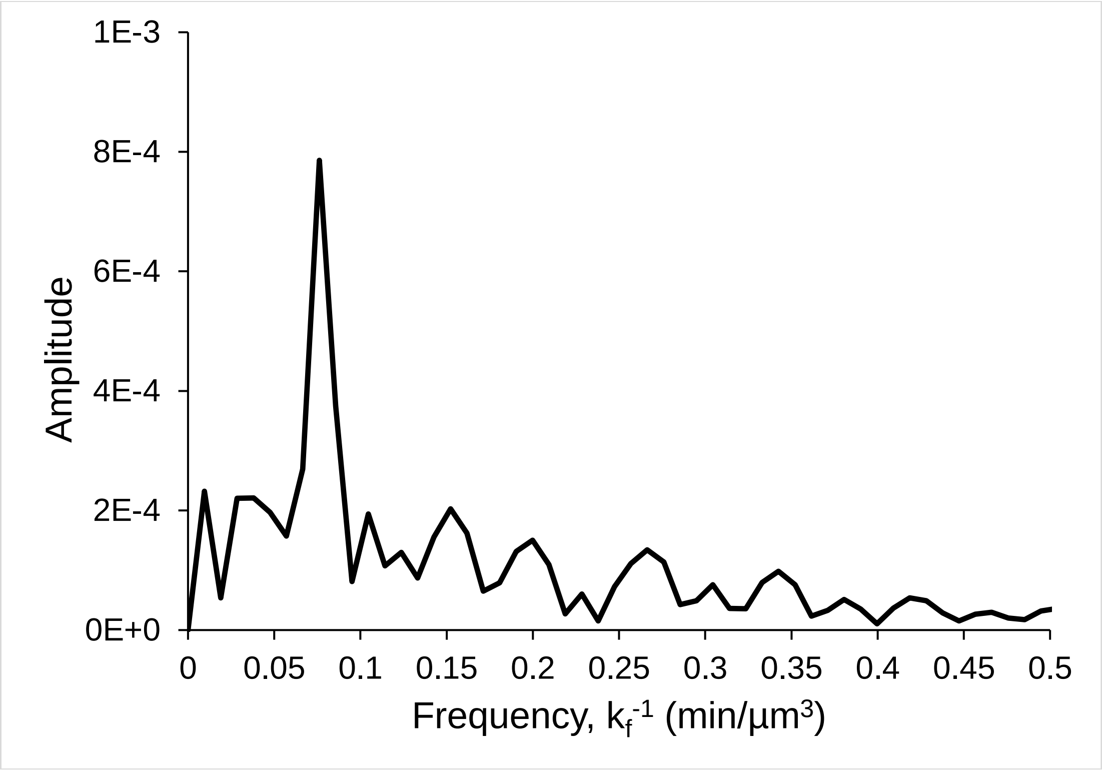


**Supplementary Figure 16. Fourier transform of the probability density distribution of influx rates of TAMRA into DNA pores**. The discrete probability density function of influx rates (*k*) in between 15 and 120 µm$^3$/min was normalized by subtracting the mean value (removing the DC bias) and removing the linear trends using the "detrend" function in MATLAB before taking the Fourier transform. The frequency peak at 0.0761 min/µm$^3$ represents a dominant periodic frequency and corresponds to a dominant period of 13.1 µm$^3$/min in the probability density function of influx rates.

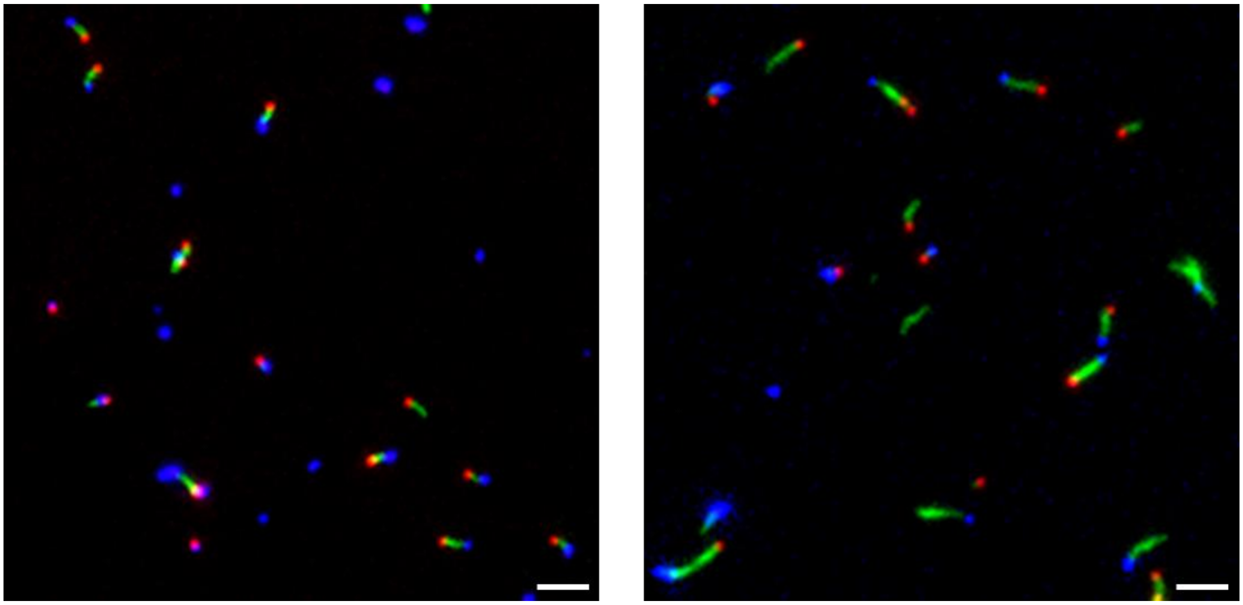

**Supplementary Figure 17. Example wide-field fluorescence image of DNA caps bound to DNA nanochannels.** The DNA nanochannels labeled with Cy3 (green) formed on pores labeled with ATTO647 (red) were mixed with two-fold concentration of caps labeled with ATTO488 (blue). The mixture was incubated at room temperature overnight before being imaged on a glass coverslip. We manually counted the nanochannels with caps at their ends in 4 images captured at random locations on the coverslip. A fraction of 0.90 ± 0.04 (95% CI, N = 244) nanochannels were capped. Scale bars, 5 μm.

**Supplementary Note 12: Computer simulation of small molecule diffusion through a DNA pore.**

First, we performed multi-resolution equilibration of the nanopore structure using the mrdna package. In the first 20 μs, the nanostructure was simulated at 1 bead per four base pair resolution, followed by an 8 μs simulation at 1 bead per nucleotide resolution. The membrane was represented as an attractive potential acting on the second turn of DNA, implicitly representing the functionalized anchors. The grid-based membrane potential (2-Å resolution in ±10 nm region in membrane plane, ±3 nm along normal axis) was generated using a custom Python script by applying a harmonic potential ($k_{\text{spring}}$ = 0.05 kcal mol$^{-1}$ Å$^{-2}$) to the distance from the surface of the toroid described by $r = 2.5 \times [1 + (1 - \text{abs}(z/2))^{-2}]$, where $z$ is the coordinate normal to the membrane for a point on the surface, and $r$ is the radius of the surface, both given in nanometers. Supplementary Movie 1 illustrates the simulation trajectory.

The microscopic configuration of the nanopore structure obtained at the end of the equilibration trajectory was used to generate grid-based representation of the membrane—nanopore system for subsequent Brownian dynamic (BD) simulations of small molecule diffusion with a fixed configuration for the pore. The unstructured single-stranded scaffold (see Supplementary Figure 4) was not included in the grid-base representation. Using such grid-based representation dramatically reduced the simulation time required for obtaining statistically significant results regarding dye molecule permeation. Each rhodamine dye was represented by a point particle that interacted with another rhodamine dye through a truncated Lennard-Jones (LJ) potential of the following form:

$$U_{dye-dye}(r) = \begin{cases} U_{LJ}(r,\epsilon) - U_{LJ}(r_{min},\epsilon) & for\ r \le r_{min} \\ 0 & for\ r > r_{min} \end{cases} \qquad (20)$$

where,

$$U_{LJ}(r,\epsilon) = 4\epsilon\left[\left(\frac{\sigma}{r}\right)^{12} - \left(\frac{\sigma}{r}\right)^{6}\right] \qquad (21)$$

was the LJ potential, $r_{min} = 2^{\frac{1}{6}}\sigma$ was the location of the potential minimum and σ = 1.2 nm and ε = 0.1 kcal/mol. Supplementary Figure 18 shows the interaction potential between the two dyes. Effectively, this potential prevented the dye molecules from forming aggregates. The DNA nanopore was represented by potential and diffusivity maps (0.5-Å and 2-Å resolution, respectively), which were generated using a custom Python script by looping over all beads in the pore in the conformation at the end of equilibration and evaluating distance-dependent functions at grid sites within the 3-nm cutoff distance of the bead. For the potential, the following function was evaluated to describe the interaction of a dye molecule with a DNA bead:

$$U_{dye-DNA}(r) = \begin{cases} U_{LJ}(r,\epsilon) - U_{LJ}(r_{min},\epsilon_{rep}) + U_{LJ}(r_{min},\epsilon_{attr}) & for\ r \le r_{min} \\ U_{LJ}(r,\epsilon_{attr}) & for\ r > r_{min} \end{cases} \qquad (22)$$

where $\epsilon_{rep} = \sqrt{0.025 \times N_{nt}}$ kcal/mol, $N_{nt}$ is the number of nucleotides represented by the DNA bead, and $\epsilon_{attr} = \alpha\epsilon_{rep}$ with σ and α being varied in different simulations to elucidate the effects of the steric and attractive interactions, respectively. The diffusivity map was generated by finding the distance at each point of the grid to the nearest DNA bead, and linearly decreasing the diffusion coefficient from the bulk value to 0.001 times the bulk value (32.2 Å$^2$/ns, obtained using the HYDROPRO server) for distances between 2 and 1 nm, mimicking the observation from all-atom MD of diminished water mobility at a distance of 1 nm from the center of a DNA duplex[11].

In addition, dye molecules were subject to a grid-based potential that confined the particles to two cylindrical regions symmetrically (18 nm radius; 78 nm height) arranged on either side of the implicitly-represented bilayer, one containing the DNA pore. The confinement potential also implicitly represented the bilayer, preventing the dye molecules from entering a 4-nm-thick slab, except in the 5-nm-radius region inside the pore. The confinement potential was created using a custom Python script that computed the distance from each voxel to the cylindrical regions described above and assigned harmonic potential with a spring constant of 1 kcal mol$^{-1}$ Å$^{-2}$. The pore was surrounded by 396 dye molecules at an approximate concentration of 4 mM all placed inside the first cylindrical region of the confinement potential that enclosed the DNA pore. The ARBD software package[11,12] was used to perform the simulations, using linear interpolation to evaluate forces from the grid-based potentials at each 50-fs timestep before BD integration was performed to advance the positions of the dye molecules. Each simulation lasted for 150 μs using a 500 fs time step. Supplementary Movie 2 illustrates a typical simulation trajectory.

To quantify the diffusion of dye molecules through the walls of the DNA nanopore, we divided the space around the DNA nanopore into three regions using three concentric cylinders centered at the cylindrical axis of DNA nanopore. The inner nanopore region was defined to be within a 2.5 nm radius of the cylindrical axis, the buffer region has a radius between 2.5 to 7 nm, and the bulk region has the radius between 7 and 12 nm. The regions were finite along the pore axis, terminating 5 nm before the pore's either end. In the beginning of each simulation, no dye molecules were present inside the nanopore. If the dye molecule entered inside the nanopore via the buffer region (within 20 ns of first entering the interior region), we counted that as a permeation event. Thus, the same dye molecules could be counted again only if it exits the nanopore volume into the bulk region and re-enters via the buffer region.

Similarly, to count the dye molecules diffusing inside the nanopore via the top cap, we divided the space surrounding the cap into three coaxial cylindrical regions: the bulk region just outside the entrance to the pore (4.5 nm radius, 7 nm height), the buffer region defined via another cylindrical region just below the bulk region and containing the entrance to the pore (4.5 nm radius, 7 nm height), and the interior of nanopore near the entrance (another cylindrical region below the buffer region having 3nm radius and 10 nm height starting from the nanopore entrance and down to the Z axis of the DNA nanopore). The bulk, buffer and interior regions are basically three cylinders kept on top of one another. In the same way as for the wall permeation calculations, the permeation event through the top end was counted if the dye molecule went from the bulk via the buffer region to the interior of the membrane region.

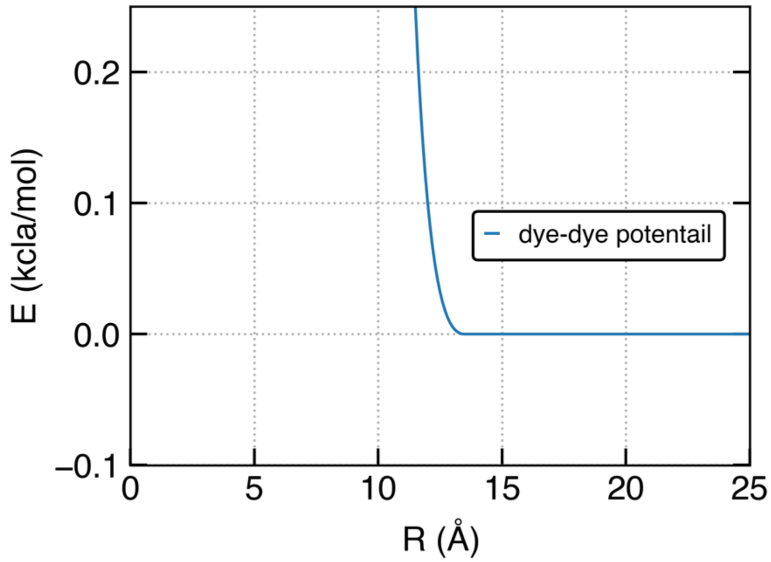


**Supplementary Figure 18. Interaction potential between two dye molecules used in the Brownian dynamics simulations of dye diffusion through the nanopore structure.** This truncated form of Lennard-Jones potential prevented aggregation of dye molecules, enabling BD simulations at high dye concentrations.

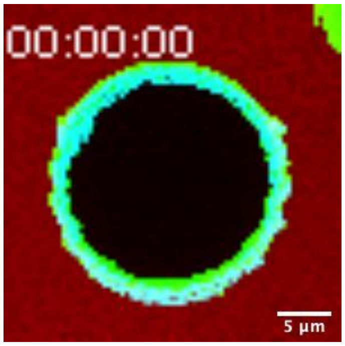


**Supplementary Movie 1. DNA nanopores induce fast TAMRA influx into a GUV.** Nanopores are labeled with ATTO647 (cyan), TAMRA in red, and GUV membranes labeled with TopFlour (green).

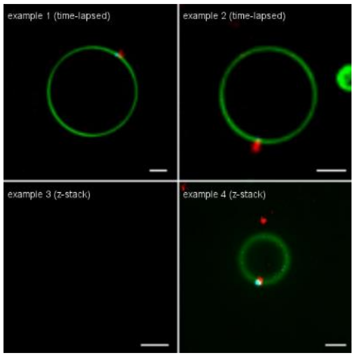


**Supplementary Movie 2. DNA nanochannels attached to DNA pores insert onto GUV membranes.** Nanochannels are labeled with Cy3 (red), nanopores labeled with ATTO647 (cyan), and GUV membranes labeled with TopFlour (green).

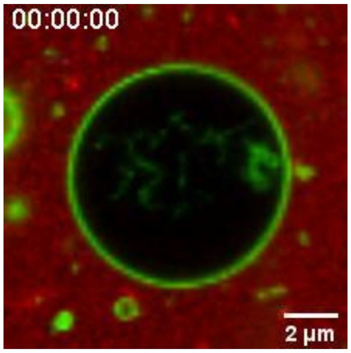


**Supplementary Movie 3. DNA nanochannels (red) induce TAMRA (red) influx into a GUV.** DNA nanochannels (Atto647, red) and TAMRA are fluorescently labeled with the same emission wavelength but nanochannels show brighter intensities in the movie. GUV membranes labeled with TopFlour (green)

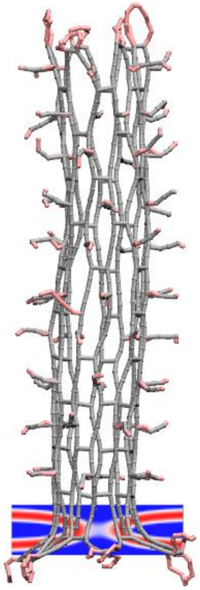

**Supplementary Movie 4. Multi-resolution equilibration of DNA pore embedded in a lipid bilayer membrane.** Starting from the caDNAno design, the DNA nanopore was simulated at 1 bead per four base pair resolution for the first 20 μs, followed by an 8 μs simulation at 1 bead per nucleotide resolution. In the first part of the movie, which corresponds to the 1 bead per 4 base pair simulation, the DNA nanostructure is represented using light grey lines. In the second part of the movie, which corresponds to the higher-resolution DNA model, the DNA nanostructure is drawn a using the VMD's Quicksurf representation, with resolution of 1 Å and the radius scale of 3.8 Å. The unfolded scaffold of the DNA origami is not shown. A planar potential applied at the bottom of the nanopore to represent the incorporation of the nanopore into the lipid bilayer membrane; the potential is shown using the RWB color map.

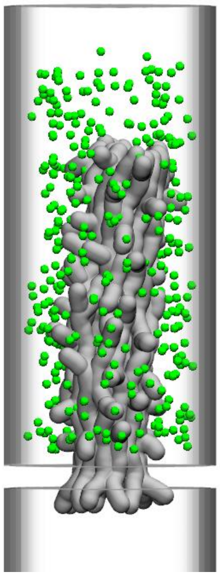

**Supplementary Movie 5. Brownian dynamics simulation of dye diffusion through the DNA pore**. The movie illustrates a representative 150 μs Brownian dynamics simulation of 396 dye molecules (green) diffusing through and around a nanopore embedded in a membrane. The interaction of DNA with the dye is described using a truncated LJ potential with σ = 16 Å. The DNA nanopore is represented using a static grid-based interaction potential and a local diffusivity map. The microscopic configuration of the DNA nanopore structure obtained at the end of the multi-resolution equilibration (Supplementary Movie 4) was used to generate this grid-based representation of the membrane-nanopore system. The DNA nanostructure is drawn using a 1 kcal $mol^{-1}$ isosurface of the dye—DNA potential map (grey). The boundary of the simulation system is shown as a semi-transparent surface.

1. Mohammed, A. M. & Schulman, R. Directing self-assembly of DNA nanotubes using programmable seeds. *Nano Letters* **13**, 4006–4013 (2013).
2. Jia, S. *et al.* Growth and site-specific organization of micron-scale biomolecular devices on living mammalian cells. (2021).
3. Li, Y. & Schulman, R. DNA Nanostructures that Self-Heal in Serum. *Nano Letters* **19**, 3751–3760 (2019).
4. Zhang, D. Y., Hariadi, R. F., Choi, H. M. T. & Winfree, E. Integrating DNA strand-displacement circuitry with DNA tile self-assembly. *Nature Communications* **4**, 1–10 (2013).
5. Douglas, S. M. *et al.* Rapid prototyping of 3D DNA-origami shapes with caDNAno. *Nucleic Acids Research* **37**, 5001–5006 (2009).
6. Maffeo, C. & Aksimentiev, A. MrDNA: a multi-resolution model for predicting the structure and dynamics of DNA systems. *Nucleic Acids Research* **48**, (2020).
7. T. E. Ouldridge et al. Structural, mechanical, and thermodynamic properties of a coarse-grained DNA model. *J. Chem. Phys*, 134, 085101 (2011).
8. Šulc, P. *et al.* Sequence-dependent thermodynamics of a coarse-grained DNA model. *Journal of Chemical Physics* **137**, 135101 (2012).

8 Octobre, G., Lemercier, C., Khochbin, S., Robert-Nicoud, M. & Souchier, C. Monitoring the interaction between DNA and a transcription factor (MEF2A) using fluorescence correlation spectroscopy. *Comptes Rendus - Biologies* **328**, 1033–1040 (2005).

9. Cheng, N. S. Formula for the viscosity of a glycerol-water mixture. *Industrial and Engineering Chemistry Research* **47**, 3285–3288 (2008).
10. Comesaña, J. F., Otero, J. J., García, E. & Correa, A. Densities and viscosities of ternary systems of water + glucose + sodium chloride at several temperatures. *Journal of Chemical and Engineering Data* **48**, 362–366 (2003).
11. Comer, J. & Aksimentiev, A. Predicting the DNA Sequence Dependence of Nanopore Ion Current Using Atomic-Resolution Brownian Dynamics. *The Journal of Physical Chemistry C* **116**, 3376–3393 (2012).
12. Ermak, D. L. & McCammon, J. A. Brownian dynamics with hydrodynamic interactions. *The Journal of Chemical Physics* **69**, 1352–1360 (1978).
